# Spatial 5mC-seq profiling of embryos and decidua after implantation in mammal

**DOI:** 10.64898/2025.12.15.694289

**Authors:** Xun Shan, Yimin Tang, Jinzhou Hu, Mingyuan Bian, Lei Gao, Jiang Liu

## Abstract

DNA methylation plays key roles in development and diseases. However, no spatial DNA methylation profiling technology has been reported until now. Here, we developed a spatial 5mC-seq method (SmC-seq) based on a microfluidic system. The SmC-seq can provide a non-biased genome-wide methylome at about single-cell scale (10 μm in width per channel). We further applied this SmC-seq to explore the spatiotemporal dynamics of DNA methylation during post-implantation development in mouse. A clear spatial heterogeneous pattern of DNA methylation among inner cell mass-derived tissues can be observed. Additionally, we identified a two-layer organization in the ectoplacental cone at the E8.5 stage, characterized by distinct DNA methylation patterning and proliferation states. Unexpectedly, a portion of maternal tissue with low DNA methylation level, enriched for nutrient-supplier progenitor cell, is observed in the middle region of maternal decidua after implantation. The hypomethylated regions in the nutrient-supplier progenitor cell cluster are associated with cell proliferation. Interestingly, the genes associated with hypomethylated regions in the mature nutrient-supplier cell cluster are enriched in exocytosis and nutrient synthesis, which is associated with nutrient provision before functional placenta is formed to support mammalian embryogenesis. In summary, SmC-seq enables spatial mapping of DNA methylation and facilitates our understanding of various biological events.

## Introduction

Spatial epigenomics technologies provide powerful tools to uncover epigenetic heterogeneity, as well as the spatial heterogeneity and organizational architecture of tissues, allowing for a more comprehensive understanding of biological systems^1–4^. DNA methylation is a well-studied epigenetic modification, which regulates gene expression, transposon silencing, and genomic stability^5,6^. Recently, single-cell DNA methylation technologies have been developed^7–9^. A limitation of these technologies is their inability to preserve the original spatial coordinates of cells, which obscures how epigenetic heterogeneity is organized and whether DNA methylation also exhibits spatial heterogeneity within tissues. In contrast, spatial DNA methylation technologies enable direct correlation of DNA methylation states with spatial locations and histological features to offer clues to the organizational principles of tissues. However, until now, no spatial DNA methylation profiling methods has been reported.

During early embryogenesis, mammalian zygotes develop into blastocysts, which consist of inner cell mass (ICM) and trophectoderm (TE). After blastocyst implantation into the mother’s uterus, both embryo and maternal uterine endometrium undergo extensive changes. In embryos, ICM differentiates into epiblast and primitive endoderm. Epiblast further differentiates into ectoderm, mesoderm and endoderm, which collectively form the fetus, while primitive endoderm differentiates into visceral endoderm (VE) and parietal endoderm^10^. Meanwhile, the TE cells consist of polar TE directly contacting with ICM, and mural TE surrounding the blastocoel. After implantation, the mural TE differentiates into primary trophoblast giant cells (TGCs), while polar TE forms the extraembryonic ectoderm (ExE) and the ectoplacental cone (EPC)^10^. The EPC appears as a triangle cap extending into the mesometrial decidua. After gastrulation, the EPC and chorionic cavities are formed, with the base of EPC cavity and extraembryonic mesoderm forming the chorion plate^10^. In mouse embryonic day 8.5 (E8.5) embryo, fusion of the chorion plate and allantois initiates hemochorial placentation^11^. Chorionic trophoblast precursors then differentiate into multinucleated syncytiotrophoblast cells through cell-cell fusion, and mononucleated sinusoid TGCs^12^. Together with endothelial cells, they constitute the maternal-fetal interface of the labyrinth zone^12,13^. As vascular networks expand, EPC-derived cells form the spongiotrophoblast layer containing spongiotrophoblast cells, glycogen trophoblast cells, and several TGC types. This layer, along with the secondary parietal-TGC (P-TGC) layer, defines the junctional zone^11,12^. The labyrinth and junctional zone together constitute the major layers of the definitive placenta that can eventually mediates nutrients and oxygen exchange between mother and fetus. Alongside embryo implantation, maternal uterine tissue undergoes decidualization to support the further development of the embryo. Extensive cell fate determinations and transitions occur during this process^11,14^.

The mammalian embryos undergo *de novo* DNA methylation after implantation^9^, which is critical for germ layer cell fate determination^15^. In contrast, TE continues to exhibit low methylation levels, which may potentially promote the invasiveness of trophoblast cells^16^. Previous studies using gene-editing cell lines and animal models have demonstrated the essential role of DNA methylation in the normal development and function of the trophoblast lineage^17,18^. For example, the loss of DNA methyltransferase Dnmt3b led to the activation of germline genes in trophoblast cells and impaired the formation of the maternal-fetal interface^19^. Furthermore, growing evidence suggests that abnormal DNA methylation is associated with the development of preeclampsia, a disorder linked to placental dysfunction^16^. Although previous single-cell DNA methylation analyses have explored embryonic tissues, such as epiblast, ectoderm, mesoderm and endoderm^9^, single-cell DNA methylation dynamics of TE during post-implantation development remain largely unexplored *in vivo*. In addition, previous study reported a stable DNA methylation status during decidualization in induced human endometrial stromal cells *in vitro*^20^. However, the dynamics of DNA methylation during decidualization *in vivo* are still unknown.

In addition, due to the lack of spatial DNA methylation method, the spatial characteristics of DNA methylation during embryonic development after implantation remain unexplored. In this study, we aim to establish a spatial 5mC-seq method and investigate the spatiotemporal dynamics of DNA methylation in embryonic and decidual cells after implantation.

## Result

### The work flow of SmC-seq

To develop a spatial 5mC-seq method (SmC-seq), we utilized a microfluidic system (Extended Data Fig. 1a) to capture DNA methylome while simultaneously preserving the spatial location information of cells in tissue (Fig. 1a). A frozen tissue section is fixed with formaldehyde to retain its structural integrity. Because genomic DNA is wrapped around histone octamers, which impedes the fragmentation for DNA methylation library preparation, tissue section was treated with three rounds of hydrochloric acid solution to remove histones from chromatin (see Methods). This step is crucial to enhance the genomic coverage of SmC-seq (Extended Data Fig. 1b). We also checked the diffusion pattern of our chip, showing no cross-channel contamination (Extended Data Fig. 1c, d). Following histone removal, Tn5 transposition is performed to fragment the genomic DNA and add adapters containing a ligation linker. Two rounds of ligation reactions are then conducted using microfluidic chips with microchannels to introduce 11-mer barcodes A and B to the DNA fragment ends. The unique combination of barcodes A and B encodes the spatial positions of cells within the tissue. As cytosine undergoes conversion to uracil during 5mC library construction, cytosine is excluded from barcodes. The DNA fragments are then reverse cross-linked and purified for subsequent DNA methylation library preparation.

**Fig. 1.**
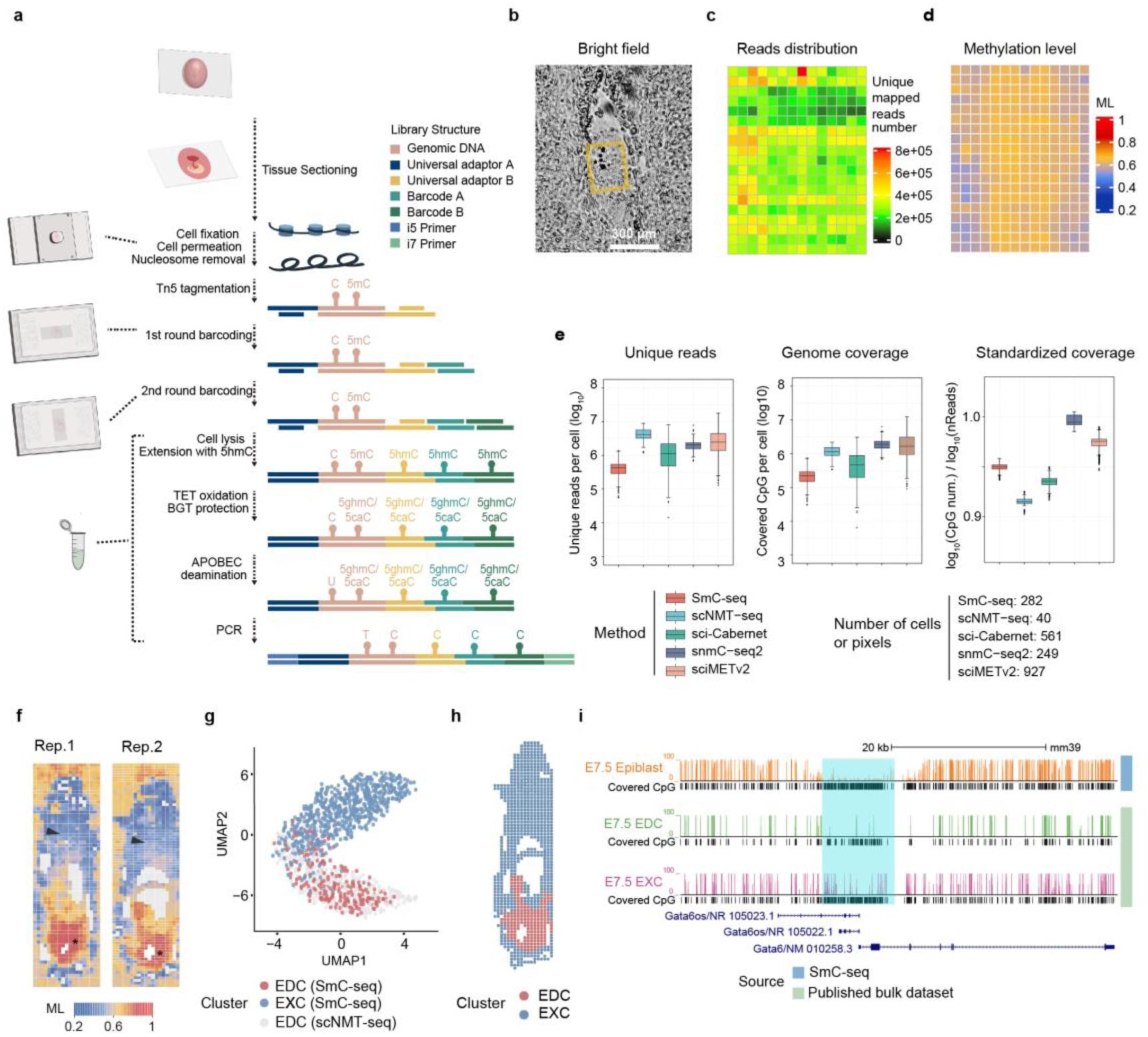
Establishment and quality assessment of spatial 5mC-seq method. **a**, The work flow of SmC-seq. **b**, Bright field image of mouse embryo-maternal tissue at the E5.5 stage. Scale bar, 300 μm. The yellow rectangle marks the embryonic region in the tissue used for SmC-seq. **c**, Spatial distribution of the number of unique mapped reads from SmC-seq data for the mouse E5.5 embryo shown in **b**. Each pixel represents a 10 µm × 10 µm region. Fluidic chips with 10-μm-width microchannels and 5-µm-interval chips were used for this analysis, with the interval walls between fluid channels omitted in the figure. **d**, Spatial distribution of 5mC levels of the mouse E5.5 embryo. Global DNA methylation level for each pixel was calculated. ML, methylation level. **e**, Box plots comparing the number of unique mapped reads (left), genome coverage (middle), and standardized genome coverage (right) among the DNA methylation sequencing data obtained using five different methods, including the SmC-seq developed in this study (10 μm × 10 μm resolution) and four published single-cell DNA methylation sequencing methods (scNMT-seq, sci-Cabernet, snmC-seq2, and sciMETv2). The numbers of pixels or cells used for comparison among these methods are shown at the bottom. Boxes and whiskers represent the 25th/75th percentiles and 1.5 x the interquartile range, respectively. **f**, Spatial distribution of 5mC levels of two replicates of mouse E7.5 embryos. Rep, replicate. Each pixel represents a 10 μm × 10 μm region. The asterisks mark the embryos, while the arrowheads mark the extraembryonic tissues. Fluidic chips with 10-μm-width microchannels and 5-μm-interval chips are used for this analysis. **g**, UMAP visualization of blastocyst-derived pixels based on the SmC-seq data from the E7.5 embryo (rep. 2) in **f**. Single-cell DNA methylation data from epiblast-derived cells generated by scNMT-seq were incorporated for comparison. The pixels from SmC-seq data were classified into two clusters, including EDC and EXC. EDC, epiblast-derived cell; EXC, extraembryonic cell. **h**, Spatial distribution of the EDC and EXC clusters in the E7.5 embryo. **i**. Genome browser view of DNA methylation levels at the *Gata6* locus in EDC and EXC clusters measured by SmC-seq, and in E7.5 embryo from published bulk whole genome bisulfite sequencing data (Li *et al*., 2023). The cyan shaded region highlights a differentially methylated region (DMR) located in the *Gata6* promoter.

Gap filling and DNA extension are performed using a dNTP mixture containing 5-hydroxy-deoxycytidine triphosphate (5-hydroxy-dCTP) instead of dCTP, which protects cytosines in the inserted sequences from deamination in the following steps. Next, an enzymatic methyl-seq procedure is employed to construct DNA methylation sequencing library^21^. During this process, TET2 and an β-glucosyltransferase (β-GT) are used to convert 5mC to 5-carboxycytosine (5caC) or glycosylated 5hmC (5ghmC), then apolipoprotein B mRNA-editing enzyme catalytic polypeptide-like (APOBEC) protein is used to deaminate unmethylated cytosines to uracils. Finally, the libraries are PCR amplified for sequencing, in which unmethylated cytosines are recognized as thymines while methylated cytosines remain as cytosines. Our method is based on EM-seq, which cannot distinguish 5mC with 5hmC. Our SmC-seq data represents a combined 5mC/5hmC signal.

To evaluate the feasibility of our SmC-seq method, we applied it to profile DNA methylation patterns in a mouse E5.5 embryo section embedded within the maternal decidual tissue (Fig. 1b). Microfluidic chips with microchannels (10μm in width per channel) were used for the experiment. The data quality was systematically assessed. The SmC-seq data show high enzymatic conversion rates, with 98.6% glycosylation protection and 98.0% deamination efficiency (Supplementary Table 1). Our data show a uniform distribution of sequencing reads across the tissue (Fig. 1c). The genome coverage of a single pixel can reach up to 3.46% (Supplementary Table 2). The average number of CpGs detected per pixel was 230,561 covering 1.05% of CpGs across the whole genome. Combining data from approximately 200 pixels achieves a genomic coverage threshold of 80% of the mouse genome (Extended Data Fig. 2a). Moreover, the sequencing reads are evenly distributed across the chromosomes and randomly distributed in different genomic elements (Extended Data Fig. 2b, c), suggesting no genomic region bias in the SmC-seq data. Analyses of our SmC data show that the global DNA methylation level of E5.5 embryos is around 0.61 (Fig. 1d and Extended Data Fig. 2d), which is consistent from the methylation level of E5.5 embryo analyzed from previous studies^9,22^ (Extended Data Fig. 2d). To benchmark the coverage of CpGs across the genome, we compared our data with four published single-cell DNA methylation sequencing datasets, generated by scNMT-seq, sci-Cabernet, snmC-seq2 and sciMETv2, respectively^7,9,23,24^. Since genome coverage positively correlates with sequencing depth, we normalized genome coverage by sequencing depth for each pixel, referred to as standardized coverage. Our results show that SmC-seq performs comparably to published single-cell methods (Fig. 1e). Scatterplot analysis of methylation signals across genomic regions at 10 kb, 100 kb, and 1 Mb resolutions show good correlation between our SmC-seq data and two public datasets (Extended Data Fig. 2e). These results suggest that SmC-seq achieves high capture efficiency for whole genome DNA methylation analysis.

To further examine the DNA methylation pattern at the pixel level detected by SmC-seq, we profiled DNA methylation in mouse E7.5 embryos and compared the data with published single-cell and bulk DNA methylation datasets of mouse embryos at the corresponding developmental stage^9,15^. The spatial patterns of DNA methylation levels in E7.5 embryo sections were consistent across two biological replicates (Fig. 1f, Extended Data Fig. 3a-c). The methylation states of about 80% of 1kb bins are similar to the previous published datasets^9,15^ (Extended Data Fig. 3d-f). These data indicate that our SmC-seq method is robust. Our data also show that DNA methylation levels are high in the pixels of epiblast-derived cells (EDCs), but low in those of extraembryonic cells (EXC) which include TE-derived cells and primitive endoderm-derived cells (Fig. 1f and Extended Data Fig. 3g). We further integrated our spatial DNA methylation data with a published single-cell DNA methylation dataset and performed clustering analysis based on DNA methylation patterns across genomic regions (see Methods). The UMAP clustering results show that these pixels can be divided into two major groups (Fig. 1g), which are annotated as the EDC cluster and the EXC cluster based on their spatial location in the mouse embryo (Fig. 1h). Consistently, the promoter methylation levels of known embryonic cell marker genes, such as *Pou5f1* and *Tdgf1*, are low in the EDC cluster but high in the EXC cluster (Extended Data Fig. 3h, i). Furthermore, the EDCs in the published single-cell data largely overlap with the annotated EDCs in our dataset, but do not overlap with the EXCs of our spatial data (Fig. 1g). We also compared the pseudo-bulk 5mC signal from our SmC-seq data with published single-cell and bulk DNA methylation datasets for mouse E7.5 embryos^9,15^, observing a good correlation (Extended Data Fig. 3j, k). A representative example is shown in Fig. 1i. In summary, the SmC-seq method is robust, and reveals a clear spatial pattern of DNA methylation in the implanted embryos.

### Embryonic and maternal uterine tissues exhibit distinct dynamic patterns of DNA methylation after implantation

To examine the spatiotemporal dynamics of DNA methylation after implantation, we collected mouse tissues containing both conceptus and maternal uterine tissues from the E6.5 to E8.5 stages to perform SmC-seq (Fig. 2a). 20-μm-wide microchannel chips were used to ensure effective coverage of both embryonic and maternal uterine tissue components for the spatial profiling of DNA methylation. Notably, DNA methylation levels vary across the tissue (Fig. 2a). Unlike the high DNA methylation levels in mature somatic cells, a significant portion of maternal decidua cells in the middle region of lateral decidua exhibit low DNA methylation levels, especially at the E7.5 stage (methylation level < 0.65) (Fig. 2a). To validate this result, we used immunofluorescent staining to examine the methylation signal in decidua. The staining results show that DNA methylation signals of some decidua cells in the middle region of lateral decidua are low at the E7.5 stage (Extended Data Fig. 4a), suggesting that DNA methylation of decidua may undergo demethylation after embryonic implantation. To exclude the possibility that low methylation level may be resulted by high proliferation, we performed co-immunostaining for 5mC and Mki67. Mki67 is a marker for replicating cells^25^. Our result show that many cells with low 5mC level show no or low Mki67 expression (Extended Data Fig. 4b). Furthermore, DNA methylation signals between Mki67^+^ and Mki67^-^ cells are comparable (Extended Data Fig. 4c). These results suggest that lowly methylated cells are not caused by cell proliferation. Our result is different from the previous report that DNA methylation remains stable during decidualization in human endometrial stromal cells under *in vitro* conditions^20^.

**Fig. 2.**
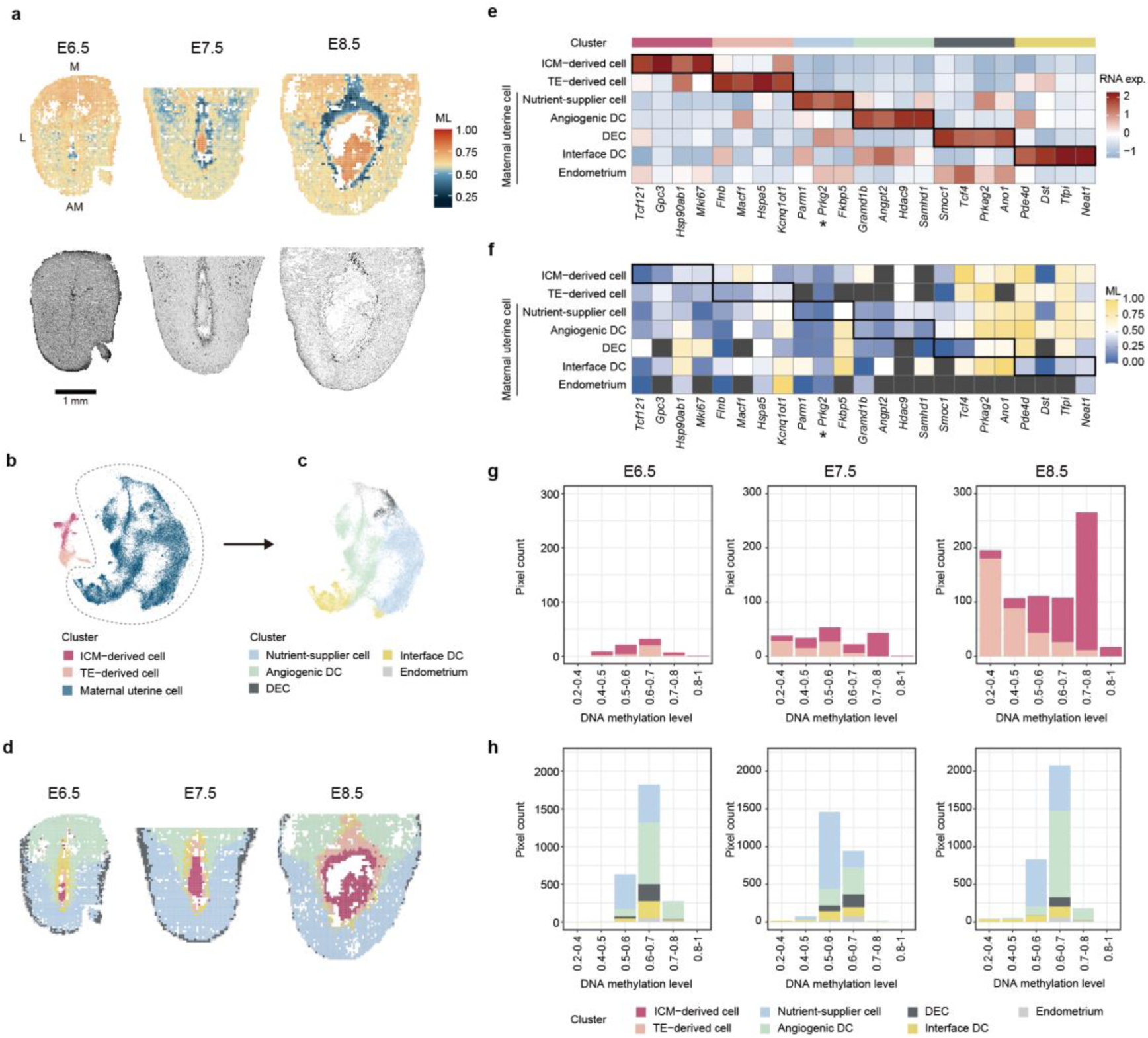
Spatial mapping and dynamics of DNA methylation levels in embryo-maternal tissues after implantation. a, Upper panels show spatial distribution of DNA methylation levels of mouse embryo-maternal tissues after implantation at the E6.5 (left), E7.5 (middle), and E8.5 stages (right). Fluidic chips with 20-μm-width microchannels were used in this analysis. Each pixel represents a 20 μm × 20 μm region, with the interval walls between fluid channels omitted in the figure. M, mesometrial; AM, antimesometrial; L, lateral; ML, methylation level. Lower panels are the bright-field image of the corresponding sections. **b-c,** UMAP visualization of clustering result based on spatial RNA-seq data from tissue sections adjacent to those used for SmC-seq in **a**. (b) Three major clusters. (c) refined clustering of the maternal uterine tissue identified in **b**. DEC, decidual endometrial cell; DC, decidual cell. **d**, Spatial distribution of different clusters in mouse embryo-maternal tissues after implantation at the E6.5 (left), E7.5 (middle), and E8.5 stages (right). Fluidic chips with 20 μm-width microchannels were used in this analysis. Each pixel represents a 20 μm × 20 μm region, with the interval walls between fluid channels omitted in the figure. The annotations of clusters are shown in **b** and **c**. **e**, Heatmap showing RNA expression levels of cell-type-specific marker genes across different clusters, classified using spatial RNA-seq data. **f,** Heatmap showing DNA methylation levels of the promoters of cell-type-specific marker genes across different clusters, based on the integrated spatial RNA-seq and SmC-seq data**. g-h,** Bar plots showing the numbers of pixels in different clusters within various DNA methylation levels ranges at different developmental stages. For example, DNA methylation levels of pixels in nutrient-supplier cell cluster are usually in the window of 0.5 – 0.6 or 0.6 – 0.7, while the methylation levels of most of pixels in TE-derived cell cluster are in the window of 0.2 – 0.4 or 0.4 – 0.5. The results of embryo-derived clusters and maternal uterine tissue-derived clusters are shown in panels **g** and **h**, respectively.

To better characterize the cellular identities, we collected the adjacent tissue slides for spatial RNA-seq analysis, assuming that neighboring tissue slides would have similar cell compositions and spatial distribution patterns. Overall, the mediam number of expressed gene of a pixel for a tissue section is around 1000 (Extended Data Fig. 4d, e). Based on the spatial RNA-seq data, all pixels can be categorized into three major clusters: ICM-derived cell cluster, TE-derived cell cluster and maternal uterine cell cluster (Fig. 2b). The marker used in identifying these clusters can be found in Table S3. Moreover, the maternal uterine cluster can be further clustered into five subclusters: endometrium, decidual endometrial cells (DECs), angiogenic decidual cells (DCs), nutrient-supplier cells, and interface DCs (Fig. 2c, d, Extended Data Fig. 4f). ICM-derived cell cluster is characterized by elevated expression levels of *Gpc3* (Fig. 2e), which encodes a membrane-bound heparin sulfate proteoglycan that functions in embryonic cell growth and differentiation^26^. TE-derived cell cluster is characterized by a trophoblast cell marker *Flnb* (Fig. 2e), which encodes Filamin B^27^. Maternal uterine cell cluster exhibits high expression level of *Prkg2* (Fig. 2e), which encodes type II cGMP-dependent protein kinase, a key regulator of decidualization^28^. The distribution of these clusters within the maternal-conceptus tissue (Fig. 1d) aligns with previous hematoxylin and eosin (H&E) staining results^29^. Furthermore, among maternal uterine cell subclusters, angiogenic DC cluster exhibits high expression of *Angpt2* (Fig. 2e), a well-known growth factor governing endothelial cells behavior and vascular remodeling^30^. Nutrient-supplier cell cluster is characterized by the active expression of several metabolism-associated genes^31^, including *Fkbp5* (also known as *Fkbp51*) (Fig. 2e).

Next, we integrated the RNA-seq-based clusters with the SmC-seq data based on tissue outlines (Extended Data Fig. 5a, b, and see Methods), and assessed DNA methylation levels across different clusters within the entire tissue. At the genome-wide level, gene expression shows negative correlation with promoter methylation level and positive correlation with gene body methylation level (Extended Data Fig. 6a, b). For example, several marker genes show a negative correlation between their expression levels and promoter methylation levels (Fig. 2e, f).

We then compared the global DNA methylation levels, defined as the average across all CpG sites detected within each pixel (see Methods), across different clusters. DNA methylation levels differ significantly among different clusters (Fig. 2g, h and Extended Data Fig. 7a). For example, DNA methylation levels of pixels in nutrient supplier cell cluster are typically in the window of 0.5–0.6 or 0.6–0.7 (Fig. 2h), while the methylation levels of most of pixels in TE-derived cell cluster are in the window of 0.2–0.4 or 0.4–0.5 (Fig. 2g).

We further investigated the dynamics of DNA methylation levels of pixels in ICM-derived cell cluster and TE-derived cell cluster during development. The DNA methylation levels of pixels in ICM-derived cell cluster gradually increase from the E6.5 to E8.5 stage (Extended Data Fig. 7b). In contrast, the average DNA methylation levels of pixels in TE-derived cell cluster decrease at the E7.5 stage and further decline at the E8.5 stage (Extended Data Fig. 7c). The decrease trend of DNA methylation in TE is further confirmed by our immunofluorescence staining (Extended Data Fig. 7d, e).

Collectively, the DNA methylation dynamics of maternal uterine and conceptus cells exhibit distinct spatial and temporal patterns at genome-wide level during post-implantation development.

### Dynamics of DNA methylation in ICM-derived tissues

Next, we explored the spatiotemporal dynamics of DNA methylation of pixels in the tissue derived from ICM from the E5.5 to E7.5 stages (Fig. 3a). The examined pixels can be categorized into three clusters based on the spatial DNA methylation data (Fig.3b and Extended Data Fig. 8a-c). Based on their position within the tissues (Fig. 3b, and Extended Data Fig. 8d) and the promoter DNA methylation levels of various cell type marker genes (Fig. 3c), these three clusters were annotated. Clusters 1, 2, and 3 are enriched for epiblast, extraembryonic endoderm (ExEndo, including visceral and parietal endoderm), and extraembryonic visceral layer (ExVL, including extraembryonic mesoderm and extraembryonic visceral endoderm), respectively. At the genome-wide level, gene expression shows negative correlation with promoter methylation level and positive correlation with gene body methylation level (Extended Data Fig. 8e, f). For example, *Ctrc* is highly methylated at its promoter and shows no expression in E7.5 embryo (Extended Data Fig. 8g). Moreover, the epiblast marker gene *Pou5f1* exhibits lower methylation at its promoter in cluster 1 (Fig. 3c), which contains an epiblast cavity^32^. Similarly, the VE marker gene *P4ha2* shows exclusive promoter hypomethylation in cluster 2 (Fig. 3c), which resides in the distal part of conceptus^33^. Yolk sac endoderm-related gene *Ttr*^34^ and yolk sac mesoderm-related gene *Cd59a*^35^ exhibit promoter hypomethylation and high expression levels in cluster 3 (Fig. 3c). At the E6.5 and E7.5 stages, the DNA methylation levels of cluster 1 which contains three germ layers do not show significant regional heterogeneity (Fig. 3a, b), indicating that the DNA methylation levels are similar among the cells of three germ layers. Consistent with previous knowledge^22^, the DNA methylation levels of the pixels in cluster 1 (epiblast) increase from the E5.5 to E6.5 stage and then remain relatively stable (Extended Data Fig. 9a).

**Fig. 3.**
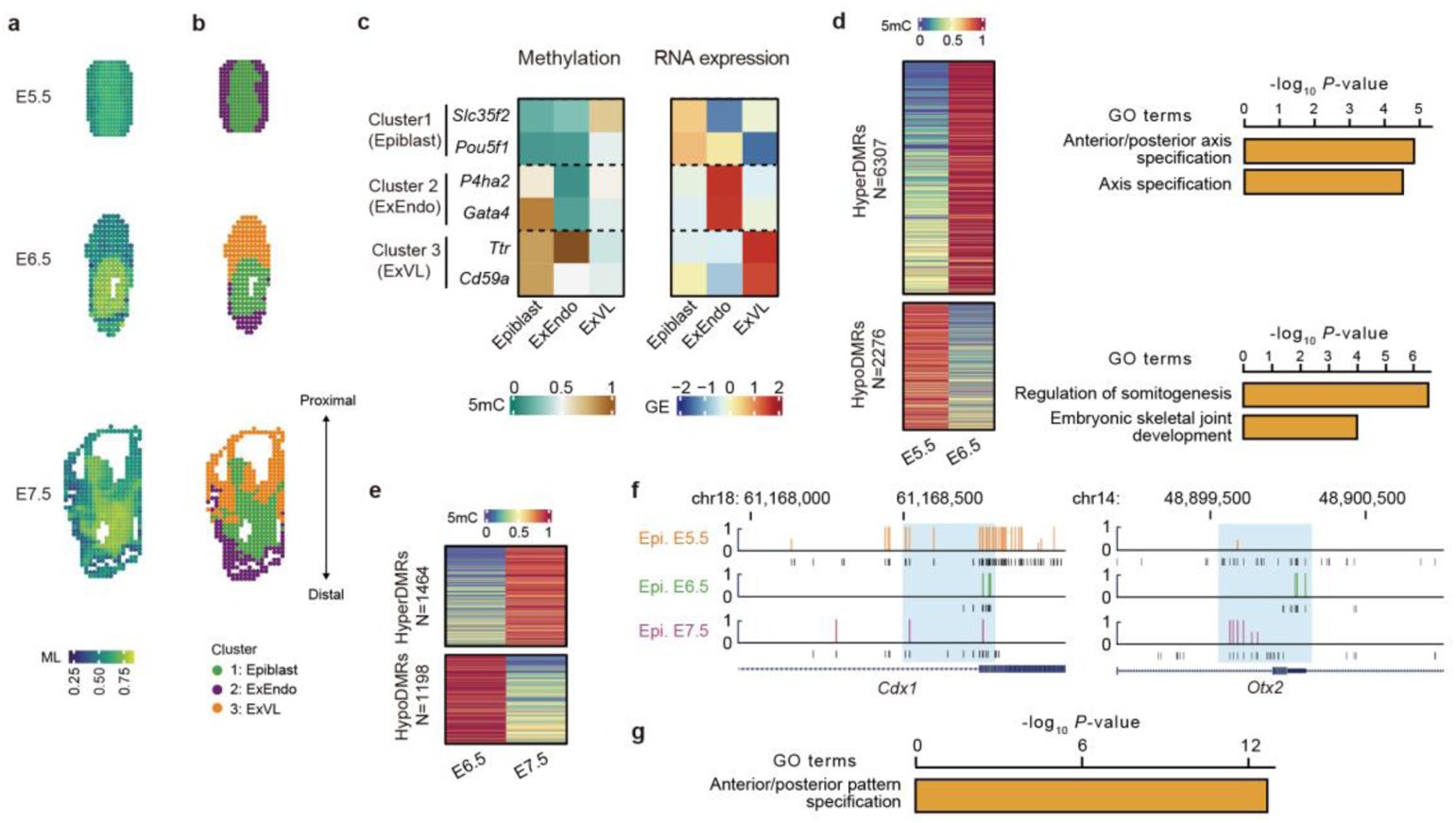
Spatial mapping and characterization of DNA methylation of ICM-derived cells in mouse embryos after implantation. **a**, Spatial distribution of DNA methylation levels of pixels within ICM-derived cells in mouse embryos at the E5.5 (top), E6.5 (middle), and E7.5 stages (bottom). Fluidic chips with 10 μm-width microchannels were used to generate these data. Each pixel represents a 10 μm × 10 μm region, with the interval walls between fluid channels omitted in the figure. ML, methylation level. 5-μm-interval chips are used for embryos at E5.5 and E7.5 stages in this analysis. **b**, Spatial distribution of clusters of ICM-derived cells in mouse embryos, based on SmC-seq data. ExEndo, extraembryonic endoderm; ExVL, extraembryonic visceral layer. **c**, Heatmap showing promoter DNA methylation levels and RNA expression levels of cell-type-specific marker genes across different clusters based on SmC-seq and spatial RNA-seq data, respectively. The clusters shown in the left part are classified based on SmC-seq data. The clusters shown in the right part are classified based on spatial RNA-seq data. 5mC, DNA methylation level of CpG sites. GE, gene expression. **d**, DNA methylation levels of DMRs in cluster 1 (epiblast) between the E5.5 and E6.5 stages, and GO enrichment of the genes associated with these DMRs. N denotes the number of DMRs. **e**, Heatmap showing DNA methylation levels of DMRs in cluster 1 (epiblast) between the E6.5 and E7.5 stages. **f**, Genome browser view of DNA methylation levels at the *Cdx1* and *Otx2* loci in cluster 1 (epiblast) at different developmental stages. Light blue shadows mark the regions of DMRs. The CpG sites covered in SmC-seq data are shown below each signal track. **g**, GO enrichment of the genes associated with hypoDMRs in cluster 2 (ExEndo) compared to cluster 3 (ExVL) at the E7.5 stage.

Then, we analyzed the differentially methylated regions (DMRs) of the cluster 1 (epiblast) at the E5.5, E6.5, and E7.5 stages. In this study, the DMRs analyses are focused on promoters and enhancers^36^. We identified 6307 hypermethylated differentially methylated regions (hyperDMRs) and 2276 hypomethylated differentially methylated regions (hypoDMRs) in E6.5 embryos compared to E5.5 embryos (Fig. 3d). 1464 hyperDMRs and 1198 hypoDMRs were identified in the E7.5 embryos compared to E6.5 embryos (Fig. 3e). We also compared the DMRs to a previously published scNMT-seq dataset of mouse early embryo during the onset of gastrulation^9^. It shows that about 28% – 47% of DMRs identified by our data are confirmed by scNMT-seq data (Extended Data Fig. 9b). Gene Ontology (GO) analyses show that genes associated with hypoDMRs at the E6.5 stage are enriched in the regulation of somitogenesis (Fig. 3d), exemplified by *Cdx1*^37^ (Fig. 3f), consistent with the determination of somitic cell fate occurring as early as the E6.5 stage. The genes associated with hyperDMRs at the E6.5 stage are enriched in anterior/posterior axis specification (Fig. 3d), exemplified by *Otx2* (Fig. 3f), consistent with that *Otx2* functions in anterior/posterior axis specification^38^. We also observed that the genes associated with hypoDMRs at the E7.5 stage, compared to the E6.5 stage, are enriched in negative regulation of cell migration (Extended Data Fig. 9c), exemplified by *Arap3*^39^ (Extended Data Fig. 9d).

Compared to epiblast cluster, DNA methylation levels are much lower in extra-embryonic tissues (clusters 2 and 3) (Extended Data Fig. 9e). These two extraembryonic clusters occupy distinct anatomical regions at the E7.5 stage. The cluster 2 are found adjacent to the distal region of conceptus, while the cluster 3 occupy the proximal region (Fig. 3b). We sought to explore the differences in DNA methylation patterns between these two clusters, as well as the potential impact of DNA methylation on development. The average DNA methylation levels are significantly higher in cluster 3 than cluster 2 (Extended Data Fig. 9e). We identified 4479 DMRs that are hypomethylated in cluster 3, and 11,405 DMRs that are hypomethylated in cluster 2. The genes associated with hypoDMRs in cluster 2 are enriched in anterior/posterior pattern specification (Fig. 3g). Given that VE is enriched in cluster 2, this result is consistent with the roles of VE in the determination of anterior-posterior polarity of embryo^40^.

### DNA methylation dynamics in maternal uterus during post-implantation development

We have classified all examined cells into three major groups: ICM-derived cells, TE-derived cells, and maternal uterine cells (Fig. 2a, b, and d). In this section, we focused on the analyses of maternal uterine cells, whose methylation dynamics remain largely unknown. To address it, we explored the spatial pattern of DNA methylation in mouse maternal uterus tissues surrounding implanted embryos from the E6.5 to E8.5 stages (Fig. 4a). The pixels of maternal uterine tissue can be classified into 9 clusters based on DNA methylation patterns (Extended Data Fig. 10a), which show distinct regional distribution within the uterine tissue (Fig. 4b). Clusters M1 and M2 show low methylation level, which are located in the lateral regions of decidua (Fig. 4a, b, and Extended Data Fig. 10b). To further annotate these clusters, we analyzed the spatial RNA-seq data of uterine cells and classified them into 9 clusters (Fig. 4c, Extended Data Fig. 10c). The figures illustrate that most of cells including nutrient-supplier-related cells belong to stromal cells (Extended Data Fig. 10c), as they express common stromal cell marker gene *vimentin*. Consistently, our staining data confirm that most of decidual cells are stromal cells (Extended Data Fig. 4a). Through image alignment of adjacent slides, we found that clusters M6 and M5 include the pixels in endometrium and decidual endometrial cell (DEC) clusters, cluster M1 is enriched for pixels in nutrient-supplier progenitor cell cluster, cluster M4 is enriched for pixels in mature nutrient-supplier cell cluster. Clusters M7 and M9 are enriched for pixels in proliferative angiogenic DSC and mature angiogenic DSC clusters, respectively. We then performed RNA velocity and pseudotime analyses for maternal decidual tissue. The results inferred a relationship between the nutrient-supplier progenitor cell cluster and mature nutrient-supplier cell cluster (Extended Data Fig. 10d, e). Interestingly, cluster M1 enriched for nutrient-supplier progenitor cell cluster exhibit the lowest DNA methylation levels than the other clusters in maternal uterine tissue, including mature nutrient-supplier cell cluster enriched in cluster M4 (Extended Data Fig. 10b). It suggests that the distinctly low DNA methylation state is a particular epigenetic state of nutrient-supplier progenitor cell cluster.

**Fig. 4.**
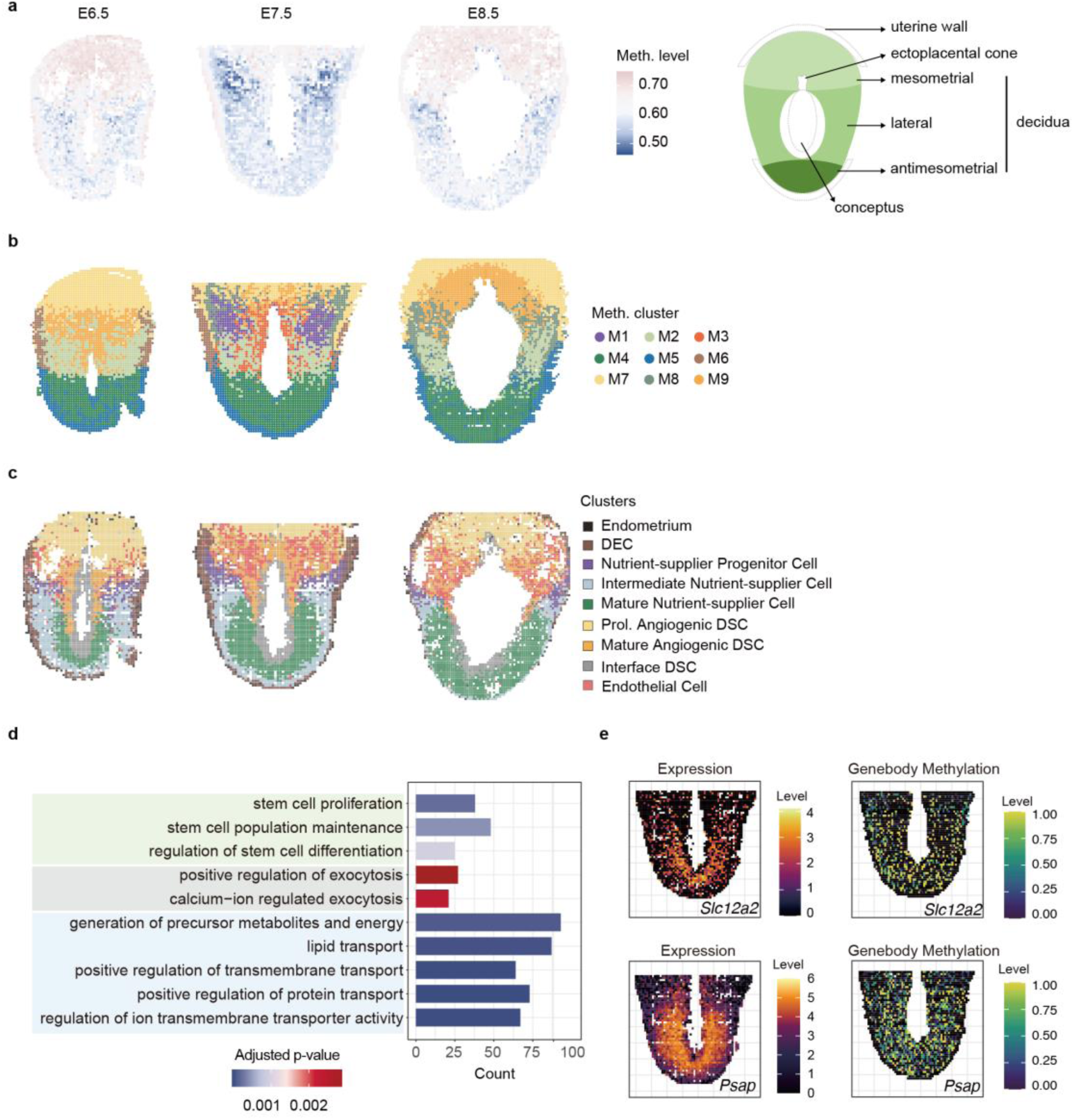
Spatial mapping and characterization of DNA methylation of maternal decidual tissue after implantation. **a**, Spatial mapping of DNA methylation levels of maternal decidual tissues at the E6.5 (left), E7.5 (middle), and E8.5 stages (right). Fluidic chips with 20 μm-width microchannels were used to generate the data. Each pixel represents a 20 μm × 20 μm region. The schematic diagram on the far right illustrates the anatomical structure of the decidua. **b**, Spatial distribution of clusters classified based on SmC-seq data of maternal decidual tissues. **c**, Spatial distribution of clusters classified based on spatial RNA-seq data of maternal decidual tissues. Fluidic chips with 20 μm-width microchannels were used to generate the data. Each pixel represents a 20 μm × 20 μm region. DEC, decidual endometrial cell; Prol, proliferative; DSC, decidual stromal cell. **d**, GO enrichment of the genes associated with the hypomethylated regions in nutrient-supplier progenitor cell cluster compared to endometrium and DEC clusters at the E7.5 stage. Those hypomethylated regions retain low DNA methylation status in mature nutrient-supplier cell cluster. **e**. Spatial distribution of RNA expression level (left) and gene body methylation level (right) of representative marker genes of mature nutrient-supplier cells in mouse maternal uterine tissue at the E7.5 stage.

To explore the potential impact of the low DNA methylation state in cluster M1, we analyzed the functions of genes associated with the hypoDMRs in cluster M1 compared with clusters M5 and M6. These hypoDMR-associated genes are enriched in pathways regulating stem cell proliferation and stem cell population maintenance (Fig. 4d), which is the reason we annotate this cluster as nutrient-supplier progenitor cell cluster. Consistently, many Mki67^+^ cells in lateral decidua exhibit low DNA methylation signal (Extended Data Fig. 4b). In addition, Slc25a33, a mitochondrial carrier protein associated with cell growth and proliferation^41^, is highly expressed in nutrient-supplier progenitor cells at the transcriptomic level (Extended Data Fig. 11a). Collectively, the data suggest that low DNA methylation state in nutrient-supplier progenitor cell is associated with cell proliferation.

Given that there is an inferred relationship between the nutrient-supplier progenitor cell cluster and mature nutrient-supplier cell cluster (Extended Data Fig. 10d, e), DNA methylation levels of pixels in cluster M4 enriched for mature nutrient-supplier cell are significantly higher than those in cluster M1 enriched for nutrient-supplier progenitor cell (Extended Data Fig. 10b). We further conducted GO analysis on genes associated with hyperDMR in cluster M4 compared to cluster M1. These genes are enriched in the processes related to somatic stem cell division and regulation of cell proliferation (Extended Data Fig. 11b). Consistently, the expression levels of genes associated with the hyperDMRs, such as a stem cell proliferation-related gene *Zbtb16*^42^, are significantly lower in mature nutrient-supplier cell cluster compared to nutrient-supplier progenitor cell cluster (Extended Data Fig. 11c). This suggests that the hypermethylation state of mature nutrient-supplier cell is associated with the attenuation of proliferative capacity. Interestingly, GO analysis of genes associated with genomic regions that maintain low DNA methylation in cluster M4 enriched for mature nutrient-supplier cells compared to cluster M1 show enrichment in positive regulation of exocytosis, regulation of generation of precursor metabolites and energy, lipid transport, and positive regulation of protein transport (Fig. 4d). Consistently, lipid transport and carboxylic acid transport are also enriched in mature nutrient-supplier cell cluster according to the result of GO analyses of genes with specific expression (Extended Data Fig. 11d). For instance, *Slc12a2* (transmembrane transport)^43^ (Fig. 4e), *Psap* (lipid metabolism)^44^ (Fig. 4e), *Glul* (nitrogen metabolism and amino acid biosynthesis)^45^ and *Fabp4* (lipid transport)^46^ (Extended Data Fig. 11e) are specifically expressed in mature nutrient-supplier cell cluster. Furthermore, Psap^+^ stromal cells are observed in the lateral decidua region, similar to the positioning of mature nutrient-supplier cell cluster near the yolk sac (Extended Data Fig. 11f). In summary, the mature nutrient-supplier cell cluster is associated with the production of various nutrients and exocytosis, suggesting their potential ability to secrete nutrients. Therefore, we designated this cluster as mature nutrient-supplier cell cluster.

Taken together, the low methylation state of nutrient-supplier progenitor cells is associated not only with their proliferative capacity but also with the nutrient provision process in mature nutrient-supplier cells.

### Dynamics of DNA methylation in TE-derived tissue

Finally, we wanted to investigate the dynamics of DNA methylation in TE-derived cells. Placenta is developed from TE. Although the potential impact of DNA methylation during early placental development has been explored, the spatiotemporal dynamics of DNA methylation during early placental development remains unclear. Here, we report the spatial DNA methylation pattern of TE-derived cells at the E8.5 stage (Fig. 5a). We performed cluster analysis based on spatial DNA methylation. Four clusters are classified (Fig. 5b, and Extended Data Fig. 12a). In contrast, only 2 clusters can be classified when using only DNA methylation data without spatial information (Extended Data Fig. 12b). Notably, clusters 1 and 2 present a clear two-layer pattern (Fig. 5b), with significantly different DNA methylation levels between these clusters (Extended Data Fig. 12c), which is confirmed by our immunofluorescence staining (Extended Data Fig. 12d-g).

**Fig. 5.**
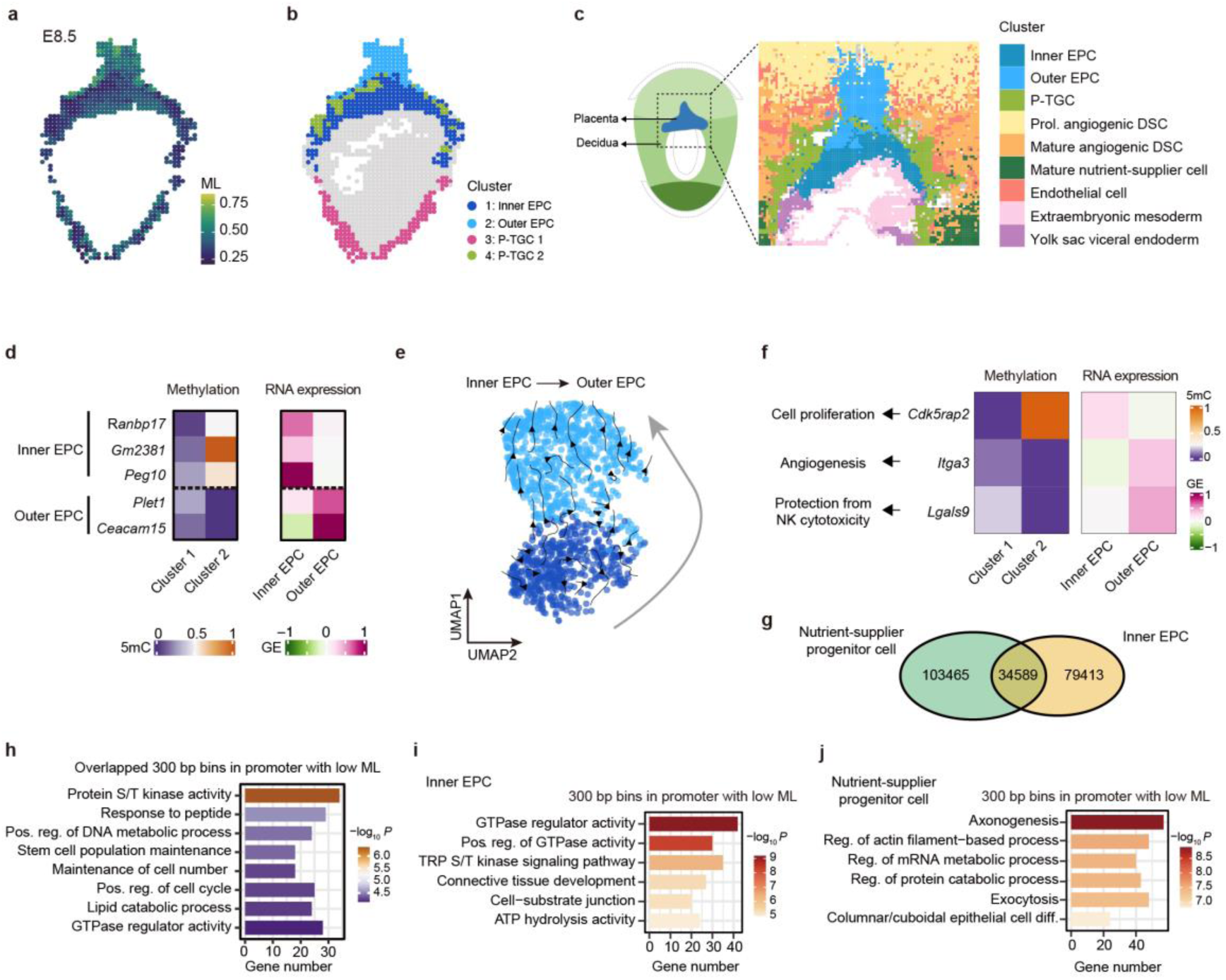
Spatial mapping and characterization of DNA methylation of trophectoderm-derived cells after implantation. a, Spatial mapping of DNA methylation levels in mouse extraembryonic tissue derived from trophectoderm at the E8.5 stage. Fluidic chips with 20 μm-width microchannels were used to generate the data. Each pixel represents a 20 μm × 20 μm region, with the interval walls between fluid channels omitted in the figure. ML, methylation level. The global DNA methylation level of each pixel is shown. **b**, Spatial distribution of clusters classified based on SmC-seq data. P-TGC, parietal-trophoblast giant cell. Prol, proliferating. **c**, Spatial distribution of clusters classified based on spatial RNA-seq data. Fluidic chips with 10 μm-width microchannels were used to generate the data. Each pixel represents a 10 μm × 10 μm region, with the interval walls between fluid channels omitted in the figure. EPC, ectoplacental cone; DSC, decidual stromal cell. The dashed rectangle in schematic diagram on the left shows the detected region within the E8.5 decidua. **d**, Heatmap showing promoter DNA methylation levels and RNA expression levels of marker genes in different clusters based on SmC-seq and spatial RNA-seq data, respectively. Clusters 1 and 2 are classified based on SmC-seq data. Clusters inner EPC and outer EPC are classified based on spatial RNA-seq data. GE, gene expression. **e**, RNA velocity analysis result of EPC. **f**, Heatmap showing promoter DNA methylation levels (left) and RNA expression levels (right) of representative genes associated with cell proliferation, angiogenesis, and protection from natural kill cell (NK) cytotoxicity, comparing inner and outer EPCs. **g**, Venn diagram showing the overlap of genomic regions (300 bp bins) with low DNA methylation levels (methylation level < 0.25) between nutrient-supplier progenitor cell cluster and inner EPC cluster. The numbers of the bins are shown. **h**, GO enrichment analysis of genes with low DNA methylation in their promoters, shared between nutrient-supplier progenitor cell cluster at the E7.5 stage and inner EPC cluster at the E8.5 stage. **i**-**j**, GO enrichment analysis of genes with promoters that are specifically lowly methylated in inner EPC cluster at the E8.5 stage (**i**) or in nutrient-supplier progenitor cell cluster at the E7.5 stage (**j**). pos, positive; reg, regulation; diff, differentiation.

To further characterize the pixel identities of TE-derived tissue, we analyzed spatial RNA-seq data of the slide which is adjacent to the one used for SmC-seq analysis. Three TE-derived clusters are identified, including the inner ectoplacental cone (EPC), outer EPC, and parietal-trophoblast giant cells (Fig. 5c and Extended Data Fig. 13a, b). We then integrated the clusters derived from SmC-seq data with those from spatial RNA-seq data. At the genome-wide level, gene expression shows negative correlation with promoter methylation level and positive correlation with gene body methylation level (Extended Data Fig. 13c, d). The promoters of inner EPC marker genes are exclusively unmethylated in cluster 1, while the promoters of outer EPC marker genes are specifically unmethylated in cluster 2 (Fig. 5d). Combined with the spatial information from DNA methylation data (Fig. 5b) and RNA data (Fig. 5c), cell clusters 1 and 2 correspond to inner EPC and outer EPC, respectively. We also observed that spatial context improved the accuracy of cell clustering based on DNA methylation data (Extended Data Fig. 13c, e).

We then sought to explore the relationship and differences between inner and outer EPC clusters. RNA velocity analysis infers that outer EPC is originated from inner EPC (Fig. 5e). The DNA methylation level of inner EPC cluster is significantly lower than that of outer EPC cluster (Fig. 5a and Extended Data Fig. 12c). We also performed GO analyses of differentially expressed genes between inner and outer EPC clusters. Our data show that cell proliferation related genes are enriched inner EPC cluster, while genes involved in angiogenesis and protection from natural killer cell mediated cytotoxicity are enriched in outer EPC cluster (Extended Data Fig. 13f, g). Additionally, cell proliferation-related genes exhibit higher expression levels in inner EPC cluster than outer EPC cluster (Extended Data Fig. 13h). Consistently, promoter DNA methylation levels of these cell proliferation-related genes are much lower in inner EPC cluster than outer EPC (Extended Data Fig. 13i), exemplified by *Cdk5rap2* (Fig. 5f). This result suggests that the inner EPC cluster is proliferative. It is confirmed by Mki67 immunostaining result (Extended Data Fig. 12d, g). Moreover, the genes involved in angiogenesis and protection from natural killer cell mediated cytotoxicity are highly expressed in outer EPC (Extended Data Fig. 13g). For example, *Igta3* and *Lgals9* have higher expression in outer EPC than inner EPC, which is consistent with the hypomethylation in *Igta3* and *Lgals9* promoters in outer EPC (Fig. 5f). We also notice that the genes whose promoters retain low DNA methylation levels in both outer and inner EPCs are enriched in the regulation of endothelial cell migration. Taken together, the DNA methylation dynamics during EPC development is associated with placental angiogenesis and immune tolerance, consistent with the role of EPCs in the invasion of conceptus cells into maternal uterine tissue and the formation of vascular system in placenta^47^.

Our data show that the DNA methylation levels of pixels for both inner EPC and nutrient-supplier progenitor cells are low, with a large number of hypomethylated regions (global methylation level < 0.25) (Fig. 5g). Some of these regions are shared by both clusters (Fig. 5g). Our results indicate that genes with hypomethylated promoters in both inner EPC and nutrient-supplier progenitor cell clusters are enriched in stem cell population maintenance and positive regulation of cell cycle (Fig. 5h). It suggests that the status of low DNA methylation is important for both EPCs and nutrient-supplier progenitor cells in maintaining their proliferative capabilities. In addition, genes with hypomethylated promoters specific to inner EPC cluster are enriched in GTPase regulator activity (Fig. 5i), whereas genes with hypomethylated promoters specific to nutrient-supplier progenitor cell cluster are enriched in exocytosis and regulation of protein catabolic processes (Fig. 5j), consistent with the previous results (Fig. 4d). Together, these findings highlight that the DNA methylation pattern of EPC and nutrient-supplier progenitor cells share certain characteristics, they also display notable differences.

In summary, a two-layer spatial DNA methylation pattern, consisting of inner EPC and outer EPC, is observed. Furthermore, DNA methylation in TE-derived tissue is associated with the formation of placenta.

## Discussion

Previously, there was no technology capable of profiling DNA methylation with spatial information. Here, we developed SmC-seq, a powerful platform for profiling the spatial DNA methylation landscape at near single-cell resolution. This method demonstrates robust performance, achieving high genome coverage of up to 3.46% of the entire genome. In this study, we employed a strategic integration of SmC-seq data with spatial RNA-seq data, enabling the precise identification of spatial dynamics of distinct cell types during development. SmC-seq method enables us to identify a previously unrecognized two-layered 5mC pattern in placenta and uncover a spatially ordered 5mC pattern in maternal decidual cells. Moreover, SmC-seq can reveal spatial heterogeneity of DNA methylation within a single cell cluster (Fig. 4). These findings offer novel insights into the spatial regulation of DNA methylation during early post-implantation development.

DNA methylation regulates gene expression and cell differentiation. However, DNA methylation states at many *cis-*regulatory elements do not always align with the differentiation state of the cell. In some cases, DNA methylation at these elements may represent a primed state that precedes cell differentiation. This may explain why the clustering patterns derived from DNA methylation and RNA expression in decidual cells differ markedly (Fig. 4b and c). This discrepancy may also be partly attributable to technical limitations, particularly the sparsity of the methylation signal, which complicates unbiased annotation and dimensionality reduction.

Before the formation of placenta, mammalian embryos rely on histotrophic absorption of essential nutrients and energy from maternal tissues. However, it remains poorly understood about how yolk sac is formed during mammalian embryogenesis. Based on our SmC-seq data, we propose that during the period of yolk sac formation, DNA demethylation occurs in the nutrient-supplier progenitor cells, which is associated with active cell proliferation. Furthermore, GO analyses show that the demethylated regions that remain in a lowly methylated state in the mature nutrient-supplier cells are enriched for nutrient-related and exocytosis-related genes. Correspondingly, the expression levels of these genes change significantly in mature nutrient-supplier cells. These findings suggest that demethylation process may reshape the epigenetic landscape, contributing to the activation of genes required for cell proliferation and the functional specialization of mature nutrient-supplier cells. Additionally, mature nutrient-supplier cells appear when yolk sac is formed, and disappear when the placenta becomes fully functional in nutrient provision to the embryo. Taken together, we propose that DNA demethylation may play a critical role in the formation of mature nutrient-supplier cells, thereby supporting nutrient production for the embryo. Additional experimental validation will be required to confirm this speculation in the future.

Our study also revealed a spatial gradient of DNA methylation within the placenta, showing a two-layered DNA methylation pattern in EPC. The two layers exhibit distinct characteristics. Inner EPCs, characterized by low global methylation levels and hypomethylation of proliferation-related genes, exhibit high proliferative capacity. In contrast, outer EPCs show higher methylation levels and hypomethylation of genes involved in angiogenesis and immune tolerance, reflecting their potential roles in placental vascularization and maternal-fetal immune interactions. These findings suggest that DNA methylation may play a critical role in the functional specialization of placental lineages.

SmC-seq enables the direct identification of regional heterogeneity within tissues. This capability is critical for unraveling the intricate spatial dynamics of DNA methylation, offering novel insights into tissue architecture and its underlying regulatory mechanisms. The versatility of this technology makes it applicable to a wide range of biological contexts and pathologies, and helps our understanding of epigenetic regulation in both health and disease.

## Methods

### Animals

This study was approved by the Ethics Committee of the Institute of Biophysics, Chinese Academy of Sciences. The experimental procedures and animal care in this study were carried out in accordance with the guidelines set by the Institutional Animal Care and Use Committee (IACUC) the Institutional Committee on Biosafety & Experimental Animal Management at the Institute of Biophysics, Chinese Academy of Sciences. Six-week-old C57BL/6N mice used in this study were purchased from the Beijing Vital River Laboratory Animal Technology Co., Ltd (Beijing, China). All animals were kept in a controlled environment with a 12-hour light/dark cycle.

### Sample collection and preparation

For the collection of embryos and uterine decidual samples, pregnant mice carrying embryos at the embryonic day 5.5 (E5.5), E6.5, E7.5, and E8.5 stage were euthanized via cervical dislocation. The uterus was collected and rinsed with ice-cold PBS solution to remove any blood. Under a microscope, the uterine endometrium was gently separated to extract the embryos, which were still wrapped in uterine decidua. The embryos were washed once more with ice-cold PBS to ensure cleanliness. They were then centered in embedding molds, and any surface fluids were removed with a pipette. Pre-cooled Tissue-Tek O.C.T. Compound (Sakura, 4583) was added over the embryos, and small bubbles were eliminated using a pipette. The molds were placed on dry ice for freezing, and the processed samples were stored at −80°C for future experiments. The frozen sample block was then sectioned into 7 μm thick slices for spatial library preparation.

### Adaptors, barcodes and other key reagents

The oligonucleotides used for SmC-seq are detailed in Supplementary Table S4, while those employed for spatial RNA sequencing (modified DBiT-seq^48^) are listed in Supplementary Table S5.

### Sample Fixation, permeabilization and histone removal

For the preparation of SmC-seq libraries, frozen tissue sections were first thawed at room temperature for 10 minutes. Next, 0.2% formaldehyde was applied to the section, and the tissue was left to react at room temperature for 10 minutes. The slide was then immersed in DEPC-treated water for 3 minutes, after which it was removed and gently blow-dried with nitrogen gas.

A mono-hole polydimethylsiloxane (PDMS) chip (Suzhou Cchip Scientific Instrument) was positioned at the center of the slide and secured with clamps. To permeabilize the tissue, 80 μL of a homemade permeabilization buffer (10 mM Tris-HCl, pH 7.4, 10 mM NaCl, 3 mM MgCl2, and 0.5% Triton-X100) was added into the hole in the chip, and the tissue was incubated at room temperature for 35 minutes. After incubation, the tissue section was washed twice with DEPC-treated water. Next, 90 μL of 0.2 N HCl was applied to the tissue section in three separate aliquots, with each addition incubated at room temperature for 15 minutes to facilitate the removal of histones from the chromatin. After the reaction, the tissue section was washed twice with DPBS to finalize the preparation.

### Labeling of spatial information on DNA fragments

To integrate adaptors into the DNA fragment flanking regions, universal adaptors A and B were first loaded onto Tn5 transposase (Novoprotein) according to the manufacturer’s instructions. The adaptors were pre-mixed in equal proportions. Then, 30 μL of the tagmentation mixture—comprising 1x homemade Tagmentation Buffer (33 mM Tris-Ac, 66 mM KAc, 10 mM MgAc_2_, 16% N,N-Dimethylformamide, 0.02% Digitonin) and 2 μL of the Tn5 transposase mixture—was applied to the tissue sample to fragment the genomic DNA and insert the adaptors. The reaction was conducted at 37°C for 45 minutes. To ensure efficient adaptor insertion, this process was repeated twice.

The mono-hole PDMS chip was substituted with a PDMS microfluidic chip containing 96 channels, each with a diameter of either 10 μm or 20 μm (Suzhou Cchip Scientific Instrument). To facilitate the ligation of the first-round barcodes to the DNA fragments, 4 μL of barcode mixture A—comprising 1x T4 Ligase Buffer (NEB), 15 U/μL T4 DNA Ligase (NEB), 0.1% Triton-X100, and 11 μM barcode A—was introduced into each channel. The mixture was then aspirated through the channels under a vacuum. The sample was subsequently incubated at 37°C for 1 hour. Following incubation, the reaction in each channel was quenched by adding 4 μL of barcode blocking mixture A (containing 4.4 μM Blocking A and 1x T4 Ligase Buffer), followed by a 20-minute incubation at 37°C. The microfluidic chip was then carefully removed, and the slide was thoroughly washed with DEPC-treated water to complete the process. To perform the ligation of the second round of barcodes, the slide was fitted with an additional PDMS microfluidic chip, oriented perpendicular to the previous one. 4 μL of barcode mixture B, consisting of 1x T4 Ligase Buffer, 15 U/μL T4 DNA Ligase, 0.1% Triton-X100, and 11 μM barcode B, was introduced into each channel. This mixture was then drawn through the channels under vacuum. The sample was incubated at 37°C for 1 hour to allow ligation. Following the incubation, the ligation reaction was quenched by adding 4 μL of barcode blocking mixture B, which contained 4.4 μM Blocking B and 1x T4 Ligase Buffer, and the reaction was further incubated at 37°C for an additional 20 minutes. After the removal of microfluidic chip, the slide was thoroughly washed with DEPC-treated water to complete the process.

After completing two rounds of barcoding, a clean mono-hole PDMS chip was positioned at the center of the slide and fixed with clamps. 50 μL of homemade DNA lysis buffer—comprising 10 mM Tris-HCl (pH 8), 5 mM EDTA, 150 mM NaCl, 0.5% SDS, and 4 mg/mL proteinase K—was added to the sample. The sample was then incubated at 55°C for 1 hour to facilitate efficient cell lysis. Following incubation, the supernatant was aspirated by pipetting 10 times and transferred to a 1.5 mL Eppendorf tube. The microfluidic hole was then rinsed twice with lysis buffer, and the collected supernatant was added to the same tube. The tube was subsequently placed in a 55°C dry bath incubator overnight to ensure complete digestion and DNA release. The DNA was then purified using SPRIselect beads (Beckman Coulter), and the final DNA was eluted with nuclease-free water for downstream applications.

### Preparation of spike-in controls

A CpG-methylated lambda phage spike-in DNA was generated by methylating the CpG sites of unmethylated lambda DNA (NEB) using CpG Methyltransferase M.SssI (NEB), following the manufacturer’s protocol. Additionally, a 5-hydroxymethylated pSP64 plasmid DNA fragment was synthesized by Takara and utilized as a 5hmC spike-in control. The CpG-methylated lambda spike-in DNA and the 5-hydroxymethylated pSP64 spike-in DNA were combined in equal ratios. The resulting spike-in DNA mixture was then tagmented using Tn5 transposase, followed by barcode ligation as previously described. The final product was stored at −20°C until further use.

### Cytosine conversion

The eluted DNA was subjected to an extension reaction using Bst DNA polymerase (NEB) to replace unmodified cytosines in the adapters and barcodes with 5-hydroxymethylated cytosines (5hmCs). The extension reaction mixture, consisting of 1x ThermoPol Buffer (NEB), 6 mM MgSO₄, 0.2 mM dATP, 0.2 mM dTTP, 0.2 mM 5-hydroxymethyl-dCTP (Abcam), 0.2 mM dGTP, 320 U/mL Bst DNA polymerase (NEB #M0275S), 0.05 ng of the spike-in DNA mix, and the eluted DNA, was incubated in a 65°C dry bath for 45 minutes. Following the reaction, the extended DNA was purified using 1.4x SPRIselect beads and eluted in nuclease-free (NF) water.

To distinguish the 5mCs in the extended DNA fragments from other cytosines, the 5mCs were oxidized by TET2, and the resulting 5hmCs or 5caC, along with the 5hmCs incorporated into the adapters and barcodes during the extension step, were glucosylated in a one-step reaction using T4 Phage β-glucosyltransferase (NEB), following the manufacturer’s instructions. The reaction mixture was incubated at 37°C in a dry bath for 2 hours. Following glucosylation, the DNA was purified using 1.4x SPRIselect beads and eluted in NF water.

For DNA denaturation, 4 µL of 0.1 M NaOH was mixed with 16 µL of the eluted DNA and incubated in a pre-heated 50°C thermal cycler for 10 minutes, with the heated lid set to 65°C. The sample was then immediately transferred to ice. The denatured DNA was deaminated using apolipoprotein B mRNA editing catalytic polypeptide (APOBEC) from the NEBNext Enzymatic Methyl-seq Conversion Module (NEB #E7125S). The reaction mixture was incubated overnight at 37°C in a thermal cycler with the heated lid set to 50°C. Finally, the DNA was purified using 1.4x SPRIselect beads and eluted in NF water.

### PCR amplification and sequencing

To set up the polymerase chain reaction (PCR), 25 μL of KAPA HiFi HotStart Uracil+ ReadyMix (2X) (KAPA), 1 μL of DNA primer A (10 μM), and 1 μL of DNA primer B (10 μM) were mixed with 23 μL of deaminated DNA and thoroughly combined. The PCR program consisted of an initial denaturation step at 95°C for 3 minutes, followed by 10 cycles of 98°C for 20 seconds, 58°C for 15 seconds, and 72°C for 60 seconds, and a final extension step at 72°C for 5 minutes. The resulting libraries were sequenced on the Illumina NovaSeq 6000 platform using paired-end 150 bp sequencing.

### Spatial transcriptome library preparation

#### Sample fixation and permeabilization

To prepare spatial transcriptome libraries, a modified version of the DBiT-seq protocol was employed. First, the frozen slide was allowed to thaw at room temperature for 10 minutes. The tissue was then fixed using 4% formaldehyde at room temperature for 20 minutes. After fixation, the slide was rinsed thoroughly with DEPC-treated water and dried using a blow-dryer. Next, the slide, a single-hole PDMS chip, and clamps were assembled following the previously described method. A 50 μL RNA permeabilization mixture, containing 0.5% Triton-X100, 0.1 U/μL RNase Inhibitor (Enzymatics), and 0.05 U/μL SUPERase In™ RNase Inhibitor (Invitrogen), was applied to the tissue and incubated at room temperature for 45 minutes. Finally, the tissue was washed twice with 50 μL of ice-cold PBS.

#### Reverse Transcription (First Strand Synthesis)

To perform first-strand cDNA synthesis, 48 μL of reverse transcription mixture was prepared, containing 12.45% PEG6000, 1x RT buffer, 25 U/μL Maxima H Minus Reverse Transcriptase (Thermo Scientific), 10 μM RT primer, 0.5 mM dNTP mix (NEB), and 0.4 U/μL RNase inhibitor (Enzymatics). This mixture was combined with 12 μL of the RNA permeabilization mixture and applied to the tissue. The slide was incubated at room temperature for 30 minutes, followed by incubation at 42°C for 90 minutes. After incubation, the slide was thoroughly washed with DEPC-treated water and blow-dried.

#### Insertion of Spatial Information into RNA

To incorporate spatial barcodes into the RNA, a PDMS microfluidic chip was assembled onto the slide as previously described. For each channel, 3 μL of RNA barcode buffer (1.8x T4 Ligase buffer, 30 U/μL T4 DNA Ligase, 0.75x NEB Buffer 3.1, 0.2% Triton-X100, 5 U/μL RNase Inhibitor (Enzymatics), and 0.1 U/μL SUPERase•In™ RNase Inhibitor) and 1 μL of RNA barcode A were added. The slide was incubated at 37°C for 1 hour. Subsequently, 4 μL of barcode blocking mixture A was added to each channel and incubated at 37°C for 20 minutes. The slide was then washed thoroughly by soaking in DEPC-treated water and blow-dried.

A second microfluidic chip, with channels oriented perpendicular to the first, was attached to the slide. For each channel, 3 μL of RNA barcode buffer and 1 μL of RNA barcode B were added, followed by incubation at 37°C for 1 hour. Next, 4 μL of barcode blocking mixture B was added to each channel and incubated at 37°C for 20 minutes. The slide was again washed thoroughly with DEPC-treated water and blow-dried. ***cDNA Purification***

A clean single-hole PDMS chip was placed onto the slide and secured with a clamp. A 50 μL aliquot of homemade RNA lysis buffer (10 mM Tris-HCl pH 8.0, 50 mM EDTA, 200 mM NaCl, 2.5% SDS, 4 mg/mL proteinase K) was added to the sample. The slide was incubated at 55°C for 2 hours. After incubation, the supernatant was pipetted 10 times to mix and transferred to a 1.5 mL tube. The hole was washed twice with lysis buffer, and the supernatant was also transferred to the same tube. The tube was stored at −80°C until further use.

For each sample, 40 μL of Dynabeads MyOne Streptavidin C1 beads (Thermo Fisher #65001) were washed three times with 1x B&W buffer (5 mM Tris-HCl pH 8.0, 0.5 mM EDTA, 1 M NaCl) supplemented with 0.05% Tween-20. The beads were then resuspended in 100 μL of 2x B&W buffer supplemented with 2 μL SUPERase•In™ RNase Inhibitor. The extracted cDNA from the stored tube was purified using the DNA Clean & Concentrator kit (Zymo #D4003) and eluted in 100 μL of NF water.

#### Reverse Transcription (Second Strand Synthesis)

The purified 100 μL DNA solution was thoroughly mixed and incubated with an equivalent volume of resuspended Dynabeads MyOne Streptavidin C1 beads at room temperature for 1 hour. The beads were then washed twice with 1x B&W buffer supplemented with 0.05% Tween-20, followed by a single wash with STE buffer (10 mM Tris pH 8.0, 1 mM EDTA, 50 mM NaCl). The beads were resuspended in a template switch mixture containing 14% PEG 6000, 1x Maxima RT buffer, 4% Ficoll PM-400, 1 mM dNTP, 2.5 μM TSO, 10 U/μL Maxima Reverse Transcriptase, and 1 U/μL RNase Inhibitor (Enzymatics). The mixture was incubated at room temperature for 30 minutes, followed by incubation at 42°C for 90 minutes. After incubation, the beads were sequentially rinsed with STE buffer and nuclease-free (NF) water.

The beads were then resuspended in PCR mixture A, consisting of 1x KAPA HiFi PCR mix (KAPA), 400 nM RNA primer A, and 400 nM RNA primer B. The cDNA was amplified using the following program: initial denaturation at 95°C for 3 minutes, followed by 6 cycles of 98°C for 30 seconds, 65°C for 45 seconds, and a final extension at 72°C for 3 minutes. The supernatant was purified using 0.8x SPRIselect beads and eluted in NF water. The DNA concentration was quantified using a Qubit 4 fluorometer (Thermo Fisher).

#### cDNA Tagmentation

The RNA Tn5 mixture was prepared by loading annealed RNA Tn5 adaptor1 and RNA Tn5 adaptor2 oligos onto Trueprep Tn5 (Vazyme) according to the manufacturer’s instructions. The tagmentation mixture, containing 1x tagment buffer L (Vazyme), 2 μL of RNA Tn5 mixture (per 50 ng cDNA), and the sample cDNA, was incubated at 55°C for 10 minutes. After tagmentation, the fragmented cDNA was purified using 0.9x SPRIselect beads and eluted in NF water.

#### PCR Amplification and Sequencing

The PCR mixture B, comprising 1x NEBNext Ultra II Q5 Master Mix (NEB), 1.25 μM RNA primer C, 1.25 μM RNA primer D, and the sample cDNA, was subjected to the following program: initial incubation at 72°C for 5 minutes, denaturation at 95°C for 3 minutes, followed by 4 cycles of 98°C for 20 seconds, 65°C for 15 seconds, and 72°C for 1 minute, with a final extension at 72°C for 2 minutes. The reaction product was purified using 0.7x SPRIselect beads and eluted in NF water. The final libraries were sequenced on the Illumina NovaSeq 6000 platform using paired-end 150 bp sequencing.

### Immunostaining

The immunostaining assays were carried out as described^49^. Briefly, the 7 μm cryosections were placed at room temperature (RT) for 10min to equilibrate. The sections were fixed with 100 μl of 2% paraformaldehyde at RT for 50 min. After two washes with 100 μl of PBS (5min each), the sections were permeabilized with 100 μl of 0.5% Triton X-100 in PBS for 40min, followed by two rounds of rinse with PBS (10min each). The slides were immersed in a Schieferdecker jar filled with 0.01 M sodium citrate pH 6.0, then placed into a microwavable dish containing pre-boiled water. The jar containing the slides was transferred to a microwave oven and heated at full power for 6min. After heating, the jar was removed and allowed to cool at RT for 10min. The slides were then rinsed in PBS in a rocking jar for 15 min. Subsequently, the sections were blocked with 100 μl of blocking solution (2% BSA, 1% goat serum, 1% donkey serum in PBS) at RT for 1 hour (hr). Primary antibodies diluted in the blocking solution (100 μl) were applied and incubated at RT for 2hrs. 5-methylcytosine antibody (Abcam, #ab10805, 1:100), Mki-67 antibody (Proteintech, #28074-1-AP, 1:200), Psap antibody (Proteintech, #10801-1-AP), Vimentin antibody (Proteintech, #10366-1-AP, 1:200), Alexa Fluor 647 anti-mouse CD31 antibody (Biolegend, #102415, 1:200), and Alexa Fluor 647 anti-Cytokeratin (pan reactive) Antibody (Biolegend, #628604, 1:200) were used in this study. After removal of the antibody solution, the slides were washed three times with 200 μL of PBST (0.2% Tween-20 in PBS, 15 min for each wash). Secondary antibodies diluted 1:250 in the blocking solution (100 μl) were applied and incubated at RT for 1hr. The secondary antibodies used in these assays included Donkey anti-mouse IgG (H+L) highly cross-adsorbed secondary antibody, Alexa Fluo 488 (Invitrogen, #A-21202), Donkey anti-rabbit IgG (H+L) highly cross-adsorbed secondary antibody, Alexa Fluor 555 (Invitrogen, #A-31572). After removal of the antibody solution, the slides were washed once with 200 μL of PBST (15 min). 100 μL of Hoechst 33342 solution (Invitrogen, #H3570, 50 μg/mL) was applied and incubated at RT for 10min. After three washes with 200 μL of PBST (15 min for each wash), the sections were mounted with cover slips using ProLong Gold Antifade Mountant (Thermo Fisher, #P36930). The images were captured using an Olympus Fluoview FV3000 confocal laser scanning microscope. Fluorescent intensities were analyzed by Image J software^50^.

### Diffusion Distance evaluation

Barcodes A DNA conjugated with fluorescent dyes was used to evaluate the diffusion distance of DNA using fluorescence microscopy. Two types of Barcodes A, labeled with Cyanine3 (Cy3) or 6-Carboxyfluorescein (6-FAM), were used. For Barcode A ligation, cells in adjacent microchannels were ligated with distinct dyes to evaluate channel leakage. The E5.5 mouse embryo frozen slide was subjected to three rounds of HCl treatment, followed by Tn5-mediated adapter ligation as described in SmC-seq procedure. A microfluidic chip featuring 10 μm-wide microchannels with 5 μm-wide intervals between them was used. 2 μM of Barcodes A labeled fluorescently were ligated to the end of Tn5 adapter at 37℃ for 1 hour. After ligation, the microchannels were washed once using Barcode blocking mixture A, and then observed under a fluorescence microscope Olympus CKX53. As described^48^, the diffusion distance is defined as the distance from the edge of microchannel to where the fluorescence signal decreased to half of its peak intensity within the microchannel. It was analyzed using Image J software.

### Spatial transcriptome data analysis

#### Quality Control, Alignment, and Normalization

During the initial processing of raw transcriptomic data, unique molecular identifiers (UMIs), RNA barcode 1, and RNA barcode 2 were extracted from the Read 2 FASTQ files. Subsequently, the Read 1 FASTQ files were aligned to the mouse reference genome (GRCm38) using STpipeline v1.7.238^51^. The aligned reads were annotated according to Gencode release M11 to generate a gene expression matrix. In this matrix, the row labels correspond to spatial coordinates derived from the combined information of RNA barcode 1 and RNA barcode 2, while the column labels represent individual genes. Next, the gene expression matrices for each sample were processed using Seurat v4.1.1^52^. Initially, quality control was conducted to filter out low-quality pixels, retaining only those with UMI counts exceeding 300 for downstream analysis. Following this, the SCTransform method was employed to independently normalize each sample. This normalization utilized a regularized negative binomial regression model, with the parameter variable.features.n=3000 set to identify and retain the top 3,000 highly variable genes for subsequent analysis.

#### Integrative Analysis of multiple datasets

To integrate our spatial transcriptomic datasets with published single-cell and spatial transcriptomic datasets, we employed Seurat’s built-in integration pipeline. Briefly, the *SelectIntegrationFeatures* function was first used to identify a set of integration features from the selected clusters, with the parameter nfeatures = 1500 to ensure a robust and representative feature set. The *PrepSCTIntegration* function was then applied to prepare the data for integration, with the anchor.features parameter set to the output of SelectIntegrationFeatures. Next, the *FindIntegrationAnchors* function was utilized to identify integration anchors, using the parameters normalization.method = “SCT” and retaining the previously defined anchor.features. Finally, data integration performed using the *IntegrateData* function, with the parameters normalization.method = “SCT” and k.weight = 50.

#### Clustering analysis and annotation

Upon completion of the integration process, the integrated dataset was designated as the default assay using the command *DefaultAssay* set to “integrated”. Principal component analysis (PCA) was subsequently performed, with the parameter npcs = 30. This was followed by UMAP dimensionality reduction, configured with the parameters reduction = “pca” and dims = 1:30 to project the data into a lower-dimensional space for visualization and interpretation. Neighboring cells were identified using the same PCA-based reduction (reduction = “pca” and dims = 1:30). Finally, clustering analysis was conducted, with the resolution parameter dynamically adjusted to reflect the specific biological context and dataset characteristics.

### GO Enrichment Analysis

The clusterProfiler package^53^ and associated databases were loaded to perform GO enrichment analysis. Gene lists were prepared and converted into Entrez gene IDs to facilitate downstream analysis. The *enrichGO* function was then applied to conduct GO enrichment analysis, with the following parameters: ont = “ALL” to include all GO categories (biological process, molecular function, and cellular component), pAdjustMethod = “BH” to correct for multiple testing using the Benjamini-Hochberg method, and pvalueCutoff = 0.01 and qvalueCutoff = 0.05 to filter for statistically significant GO terms. Finally, the enrichment results were visualized using the ggplot2 package, generating bar plots and bubble plots to effectively display the significantly enriched GO terms.

### Cell Differentiation Trajectory Analysis

To elucidate the dynamic changes and developmental processes underlying cell lineages, two complementary methods were employed for inferring cell differentiation trajectories: RNA velocity-based inference and pseudotime analysis.

RNA velocity analysis was performed using scVelo v0.2.4^54^, leveraging RNA splicing information to infer developmental trajectories. The abundances of spliced and unspliced mRNAs were quantified using Velocyto v0.17.17^55^ with default parameters. k-nearest neighbor graph (k=8) was constructed, with redundant meta-cells (n>5) removed to streamline the analysis. The count matrix was then filtered and normalized using the *scv.pp.filter_and_normalize* function in scVelo. RNA synthesis and degradation rates were estimated using a kinetic model implemented in the *scv.tl.recover_dynamics* function, and velocity vectors were computed based on cosine similarity between meta-cells, capturing the direction and magnitude of transcriptional changes. Finally, UMAP embeddings derived from Seurat analysis were integrated with RNA velocity data, and the *scv.pl.velocity_embedding_stream* function was used to visualize RNA velocities as streamlines.

### SmC-seq data analysis

The analysis of spatial methylation data involves several key steps, which have been integrated into an analysis pipeline. The integrated pipeline can directly process raw FASTQ files and generate outputs containing both spatial information and DNA methylation profile for the corresponding tissue section.

### Quality control

To ensure the quality of sequencing data from the methylation library, a quality control step was implemented, in which reads with unmatched barcode combinations were removed using UMI-tools (version 1.1.2)^56^. Crick and Watson strand identities were then determined based on specific bases located between two rounds of barcodes and processed separately in the subsequent analysis. Finally, adapter removal and low-quality read filtering were carried out using Trim Galore (version 0.6.6).

### Sequence read alignment

For spatial DNA methylation raw data, DNA barcode A and B sequences were extracted from Read 2 of the FASTQ file. Filtered reads were then mapped to the mouse genome (GRCm39)^57^, followed by an iterative mapping step to improve the mapping rate. This step included trimming unmapped reads by 5 bp at their 5’ end and re-mapping them to the mouse genome, which was repeated five times. Subsequently, the resulting files from the iterative mapping process were merged to obtain the final optimal result. The BAM file was then divided into multiple subsets based on the unique combinations of two rounds of barcodes, with each subset containing only a single barcode combination representing a distinct pixel.

### Deduplication and methylation level calculation

The BAM file subsets were subjected to deduplication using the *deduplicate_bismark* function in Bismark (v.0.23.0). Subsequently, the DNA methylation profile for each subset was extracted using *methylation_extractor* function in Bismark. The spatial distribution of methylation levels was then calculated and visualized using ComplexHeatmap (v.2.14.0)^58^.

### Pixel alignment

The integration of DNA methylation and RNA spatial data was achieved through the spatial alignment of their respective images using an affine transformation. Initially, both the SmC-seq methylation data and DBiT-seq RNA images were preprocessed by converting them to grayscale and downsampling to 192×192 pixels, preserving spatial structure while reducing resolution^59^. Binary edge detection was applied to highlight tissue boundaries, aiding in alignment. Affine image registration was performed using the SimpleITK package^60^, with the RNA image as the fixed image and the methylation image as the moving image. The registration process involved manual tuning steps, including grayscale threshold adjustment, Gaussian smoothing, and affine transformation optimization, which were iterated for best alignment. The quality of the alignment was assessed using the Dice coefficient, ensuring a strong spatial overlap (Dice coefficient > 0.85) between the methylation and RNA dotplots. The resulting affine transformation matrix was applied to the methylation data to map it onto the RNA spatial framework. Finally, a filtering step was implemented to exclude spatially unreliable methylation points (those shifting more than 10 μm from the nearest RNA pixel), ensuring only the most accurate data were retained for joint analysis of RNA expression and DNA methylation patterns.

### Co-clustering analysis

To validate the quality of the spatial methylation library, we conducted unsupervised co-clustering of our SmC-seq data with previously published single-cell methylation data from mouse embryos^9^. DNA methylation levels of all gene body regions after quality control were used as features for co-clustering analysis. The average DNA methylation level was calculated by the formula: Average DNA methylation levels = Number of CpG with methylation / (Number of CpG with methylation + Number of CpG without methylation). These regions were filtered based on the criterion that more than 5 CpGs were covered in over 20% of all pixels. A DNA methylation matrix was generated, with columns representing features and rows representing individual pixels in SmC-seq data or cells in the published single-cell-5mC data. Empty values in the matrix were replaced with the average of non-empty values in the same column. Our data also show that DNA methylation levels of genebody or promoter regions are usually 0 and 1. Because the average methylation level is close to 0.5 and the probability of 0.5 is relatively low, we filled the missing values with 0.5 for one column with excessive missing values, which will have limited effects on the features of the matrix. The top 20 principal components (PCs) for each pixel or cell were computed using the prcomp function in the R package stats (v.4.2.3). Finally, after dimension reduction via UMAP, the results were visualized in 2D using the R package umap (v.0.2.10.0).

### Clustering analysis of spatial DNA methylation data

To simultaneously incorporate spatial locations and methylation levels in clustering analysis, we implemented a novel clustering method, which integrated the Euclidean distance matrix with the methylation level matrix. We first generated a Euclidean distance matrix for all pixels (e.g., for a slice with 96 pixels*96 pixels, a matrix with 9216 rows and 9216 columns). Then we identified the critical anchors with important spatial information by using the FindIntegrationAnchors function in Seurat v4.1.1. Next, the weight of the dimensionally reduced spatial matrix and the methylation level matrix is the same in the downstream analysis. This was followed by the principal component analysis and clustering using the K-means algorithm from the R package stats (v.4.2.3). We also combined k-means with the get_clust_tendency and clusGap functions in the R package factoextra (v.1.0.7) to identify the optimal cluster number. Finally, the selected PCs were dimensionally reduced into a 2D object using the UMAP algorithm in the R package umap. The in situ spatial distribution of all these clusters was visualized.

### DMR analysis and functional enrichment analysis

To perform DMR analysis, the genome was divided into 300 bp bins, which were filtered to include only those containing more than five CpGs covered by sequencing data. Significance was assessed using Fisher’s extract test in the R package stats (v.4.2.3). DMRs were selected based on a p-value threshold of < 0.01. To control the False Discovery Rate (FDR) in multiple hypothesis testing of DMRs, we performed the Benjamini-Hochberg (BH) method, and the cutoff level of FDR is 0.1. We identified DMR-associated promoters, defining the promoter region as the 2 kb upstream and 500 bp downstream of the transcription start site (TSS) for each gene. We also identified DMR-associated genes, defined as the gene closest to the DMR. GO term enrichment analyses for both DMR-associated promoters and genes were conducted through a series of R packages, including clusterProfiler (v.4.6.2)^53^, enrichplot (v.1.18.4), org.Mm.eg.db (v.3.16.0), and biomaRt (v.2.54.1)^61^.

## Statistical analysis

Statistical analysis was performed using R (v.4.2.3). The statistical tests used in this study were specified in figure legends or bioinformatics analysis sections.

## Data availability

The raw sequence data of SmC-seq and spatial RNA-seq reported in this paper have been deposited in the Genome Sequence Archive in National Genomics Data Center China National Center for Bioinformation

(Accession number: CRA023723, https://ngdc.cncb.ac.cn/gsa/s/Fmjsf7Jv; CRA028148, https://ngdc.cncb.ac.cn/gsa/s/X8lEEn6P).

## Code availability

The code used in this study is deposited in github (https://github.com/LiuLab888/spatial-5mC-project/).

## Acknowledgements

This study was supported by the grants from National Key Research and Development Program of China (2024YFA1802100 to L.G.), National Natural Science Foundation of China (32430024 to J.L), National Key Research and Development Program of China (2024YFA0916603 to J.L., 2022YFC2703302 to L.G.), National Natural Science Foundation of China (32471502 to L.G.), and Strategic Priority Research Program of the Chinese Academy of Sciences (XDB1000302 to J.L.). We thank Shuqin Jia for assistance with reagent ordering and administrative support, , and Zhuanzhuan Xing for help with the animal experiments.

## Author contributions

J.L. supervised the project; J.L. and L.G. conceptualized the project; X.S., L.G., and Y.T. designed and developed the method; X.S., J.H., and L.G. performed the experiments; Y.T., X.S., J.H., and L.G. performed computational analysis; X.S., L.G., Y.T. and J.L. wrote the original draft; All authors contributed to revising the manuscript and approved the final version.

## Competing interests

Authors declare that they have no competing interests.

## Materials & Correspondence

Correspondence and requests for materials should be addressed to Jiang Liu and Lei Gao

---

**Extended Data Fig.1.**
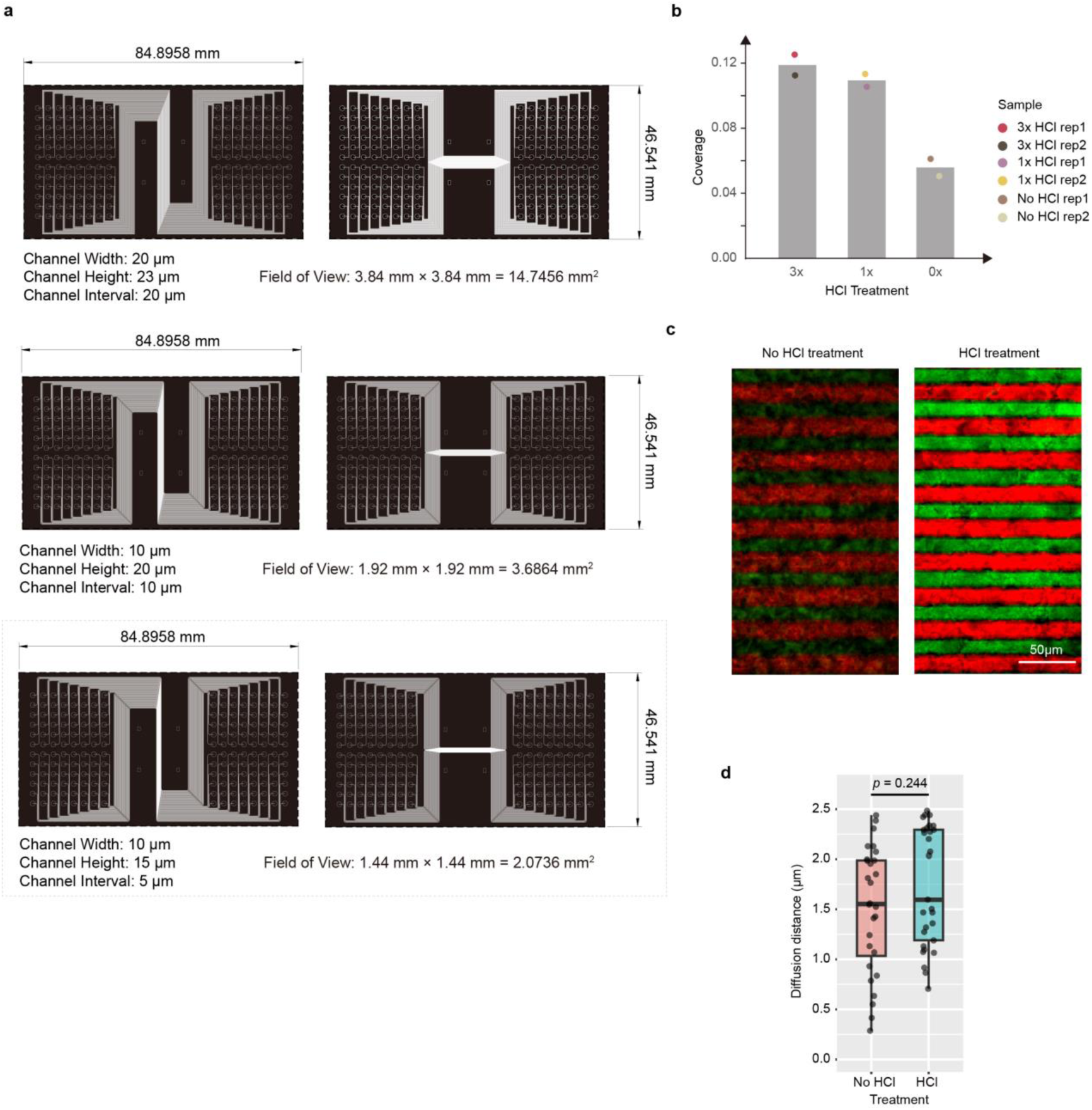
Microchip design and effects of HCl treatment on DNA diffusion. **a**, Schematic designs of the three versions of microchannel chips used in this study. **b**, Genome coverage of SmC-seq at the pseudobulk level under three different HCl treatment conditions. “3× HCl” indicates treatment with 0.2 N HCl at room temperature for 5 min per round, repeated for three rounds; “1× HCl” indicates a single round of the same treatment; “No HCl” indicates no HCl treatment. “Rep” denotes replicate. For each of the six samples, 69,256,892 raw reads were used as input. Genome coverage was defined as the proportion of CpGs across the whole genome that were covered by sequencing reads. **c**, Fluorescent imaging measuring cross-channel diffusion distances and possible crosstalk between two neighboring channels by alternately flowing 5-Carboxyfluorescein (5-FAM, Green)-labeled Barcode A and Cyanines3 (Cy3, Red)-labeled Barcode A in adjacent channels. E5.5 embryonic tissue sections, with or without three-rounds of HCl treatment, were evaluated and compared. The microchannel chip used here consists of 10 μm width channels with 5 μm spacing. **d**, Quantification of diffusion distances with and without three-rounds of HCl treatment. Two-side t-test was used for statistical analysis.

**Extended Data Fig.2.**
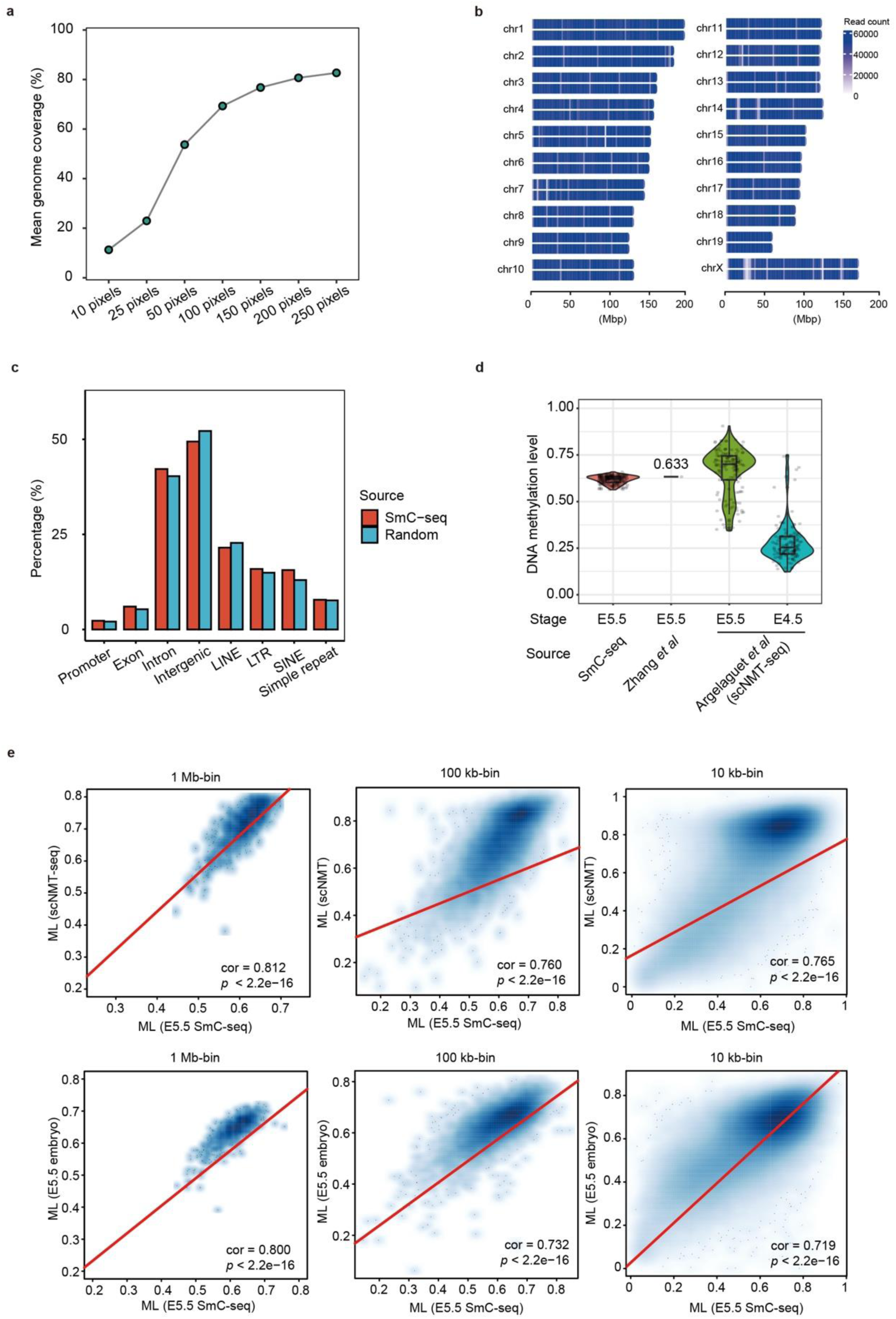
Evaluation SmC-seq in E5.5 mouse embryo. a, Line chart showing CpG genome coverage based on pseudo-bulk data generated by merging different number of pixels (10 μm × 10 μm) in the SmC-seq data of the E5.5 embryo. **b,** Read distribution from the SmC-seq data of the E5.5 embryo across all chromosomes in the mouse. The number of reads in each non-overlapping 1-million-base-pair (Mbp) bin is shown. For each chromosome, the upper strand represents the Watson strand, while the bottom strand represents the Crick strand. **c,** Bar chart depicting the percentages of different genome elements covered by E5.5 mouse embryo sequencing data using SmC-seq (red) compared with a set of random genomic regions matched for sequencing read length and number (blue). **d,** DNA methylation levels of mouse E5.5 epiblast (embryo) measured by SmC-seq or from published data. The published data including bulk DNA methylation sequencing data and scNMT-seq data, were obtained from Zhang *et al*., (2018) and Argelaguet *et al*., (2019), respectively. The global DNA methylation level was calculated for each cell or sample. **e,** Scatter plots comparing DNA methylation levels in non-overlapping bins of various sizes (1 Mb, 100 kb, 10 kb) between E5.5 embryo measured by SmC-seq and E5.5 epiblast cells measured by scNMT-seq (upper) or bulk DNA methylation sequencing (bottom). Pearson’s correlation coefficients were shown (cor). Linear regression lines are included.

**Extended Data Fig.3.**
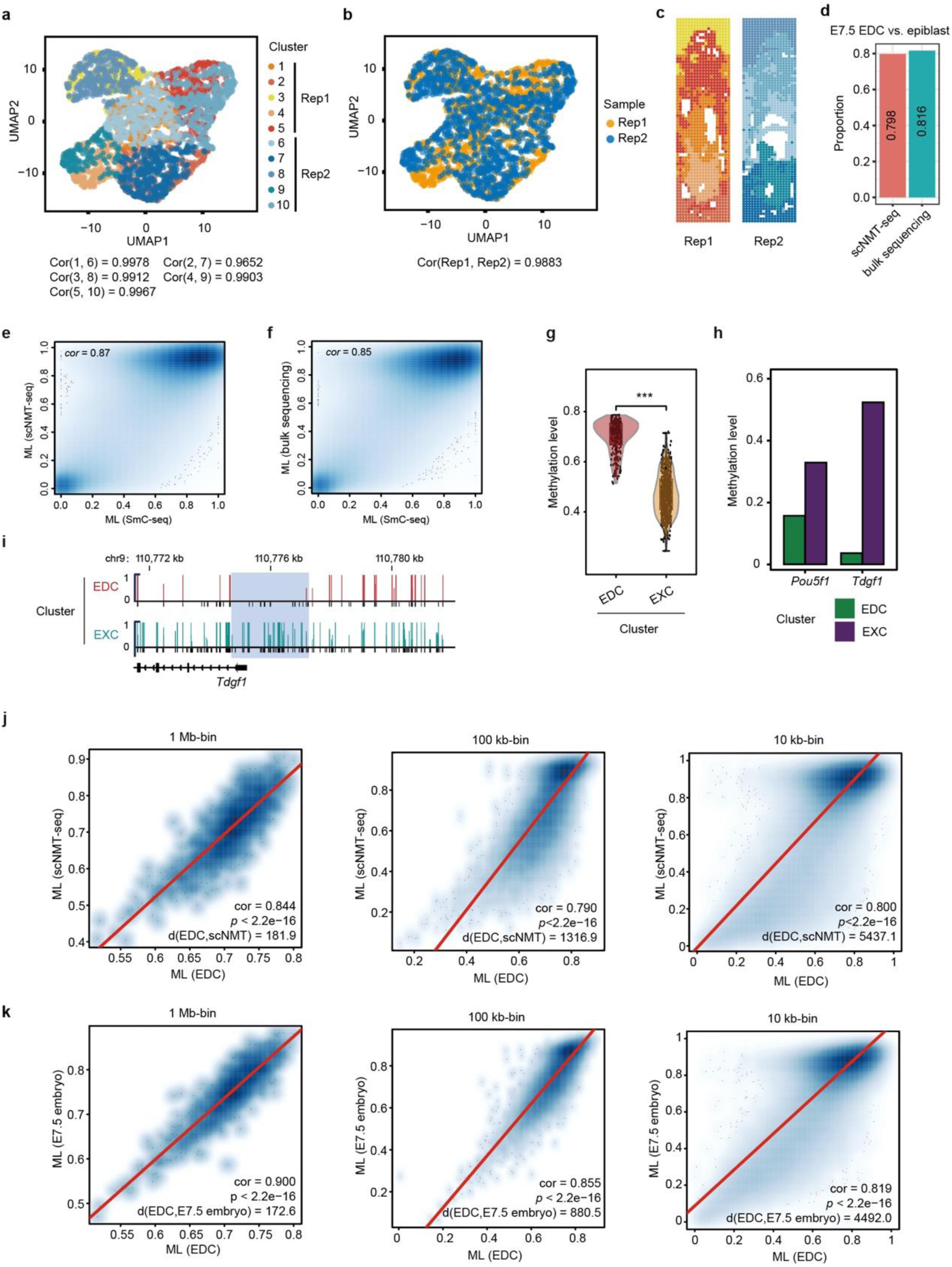
Evaluation SmC-seq in E7.5 mouse embryo. a-b, UMAP visualization of SmC-seq data from two E7.5 embryo replicates, each at a 10 µm × 10 µm resolution, colored by clusters (**a**) or samples (**b**). The correlation coefficients are computed based on the global methylation levels of pixels for all overlapping cluster pairs in the UMAP plot (**a**) from different replicates. For example, Cor(3, 8) denotes the correlation coefficients between cluster 3 in replicate 1 and cluster 8 in replicate 2. The overall correlation coefficient between replicates (**b**) is then calculated as the mean of all such pairwise correlations. Cor, correlation coefficient; Rep, replicate. **c,** Spatial localization of distinct clusters (presented in **a**) within the corresponding E7.5 embryo section replicates. The colors refer to the clusters annotated in **a**. **d,** Proportion of 1kb bins showing consistent methylation levels between SmC-seq data and other published data. The published data were obtained from Li *et al*., (2023) and Argelaguet *et al*., (2019). Bins that do not meet the criteria (BH-adjusted *p*-value < 0.05 and methylation level difference ≥ 0.2) are considered to have consistent methylation levels. **e-f,** Scatter plot comparing DNA methylation levels of non-overlapping 1kb bins between EDC cluster from SmC-seq data and epiblast from public scNMT-seq data (**e**) or bulk sequencing data (**f**). **g,** Comparison of DNA methylation levels between EDC and EXC clusters in mouse E7.5 embryo. Two-sided wilcoxon rank sum test was performed. *** represents *p-*value < 0.001. EDC, epiblast-derived cell; EXC, extraembryonic cell. **h,** DNA methylation levels of the promoters of well-known embryonic cell marker genes in EDC and EXC clusters. **i,** Genome browser view of pseudo-bulk DNA methylation levels at the *Tdgf1* promoter in EDC and EXC clusters. Light blue shadow marks the promoter of *Tdgf1*. The covered CpG sites in pseudo-bulk DNA methylation data are indicated below each track. **j.** Scatter plots comparing DNA methylation levels in non-overlapping bins of various sizes (1Mb, 100kb, 10kb) between EDC cluster at the E7.5 stage measured by SmC-seq and E7.5 epiblast cells measured by scNMT-seq (Argelaguet *et al*., 2019). Pearson’s correlation coeffiecients (cor) and Euclidean distances (d) are shown. Linear regression lines are included. **k,** Scatter plots comparing DNA methylation levels in non-overlapping bins of various sizes (1Mb, 100kb, 10kb) between EDC cluster measured by SmC-seq and E7.5 epiblast cells measured by bulk whole genome bisulfite sequencing (Li *et al*., 2023).

**Extended Data Fig.4.**
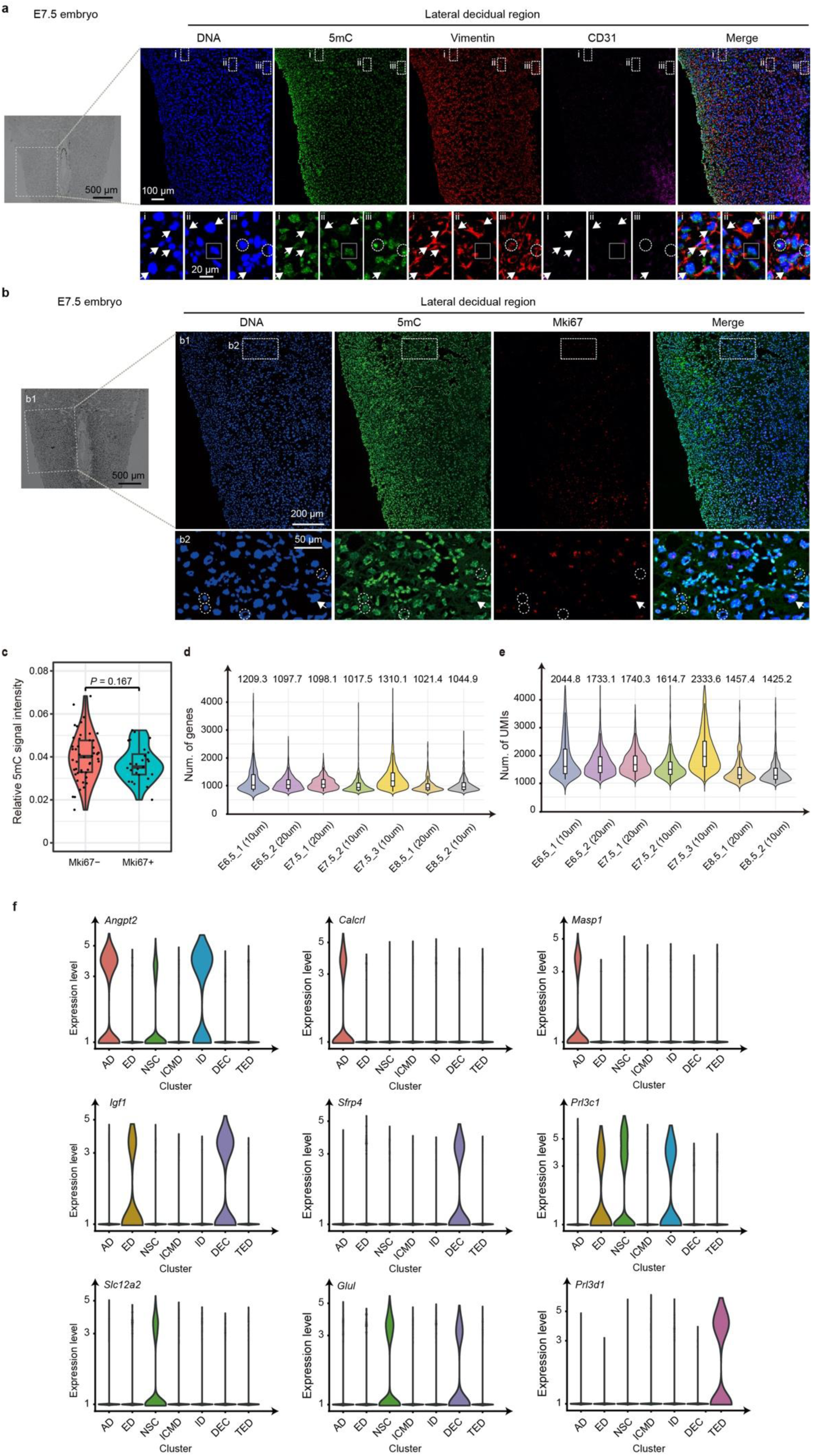
Immunostaining and spatial transcriptomic profiling of E7.5 decidua. a, Immunostaining for 5mC, Vimentin and Cd31 in the lateral decidua at the E7.5 stage. Vimentin is a marker of stromal cell. Cd31 is a marker of endothelial cell. Arrows indicate stromal cells with low 5mC signal. Dashed circles indicate stromal cells with strong 5mC signal. Solid rectangles indicate a non-stromal cell with strong 5mC signal. Magnified views of regions i, ii, and iii are shown at the bottom. The data show that most of cells have positive staining of Vimentin, in contrast a small proportion of cells have positive staining of Cd31. **b,** Immunostaining for 5mC and Mki67 in the lateral decidua at the E7.5 stage. Arrows indicate stromal cells with low 5mC signal. Dashed circles indicate stromal cells with strong 5mC signal. Magnified views of the regions within the dashed rectangles are shown. **c,** Comparison of relative 5mC fluorescent signal intensities between Mki67^+^ and Mki67^-^ cells. The analysis was performed on the cells shown in the zoomed-in region in panel **b**. Wilcoxon rank-sum test was used. **d,** Violin plots showing the number of genes detected per pixel across different samples in the spatial transcriptomic sequencing data. The widths of microchannel in fluidic chips used to generate corresponding data are indicated in parentheses. **e,** Violin plots showing the number of unique molecular identifiers (UMIs) detected per pixel across different samples. **f,** Violin plots displaying the expression levels of representative marker genes across major clusters in embryo–uterus tissue, as detected by spatial transcriptomic sequencing. AD, angiogenic decidual cell; ED, endometrium; NSC, nutrient-supplier cell; ICMD, ICM-derived cell; ID, interface decidual cell; DEC, decidual endometrial cell; TED, TE-derived cell.

**Extended Data Fig.5.**
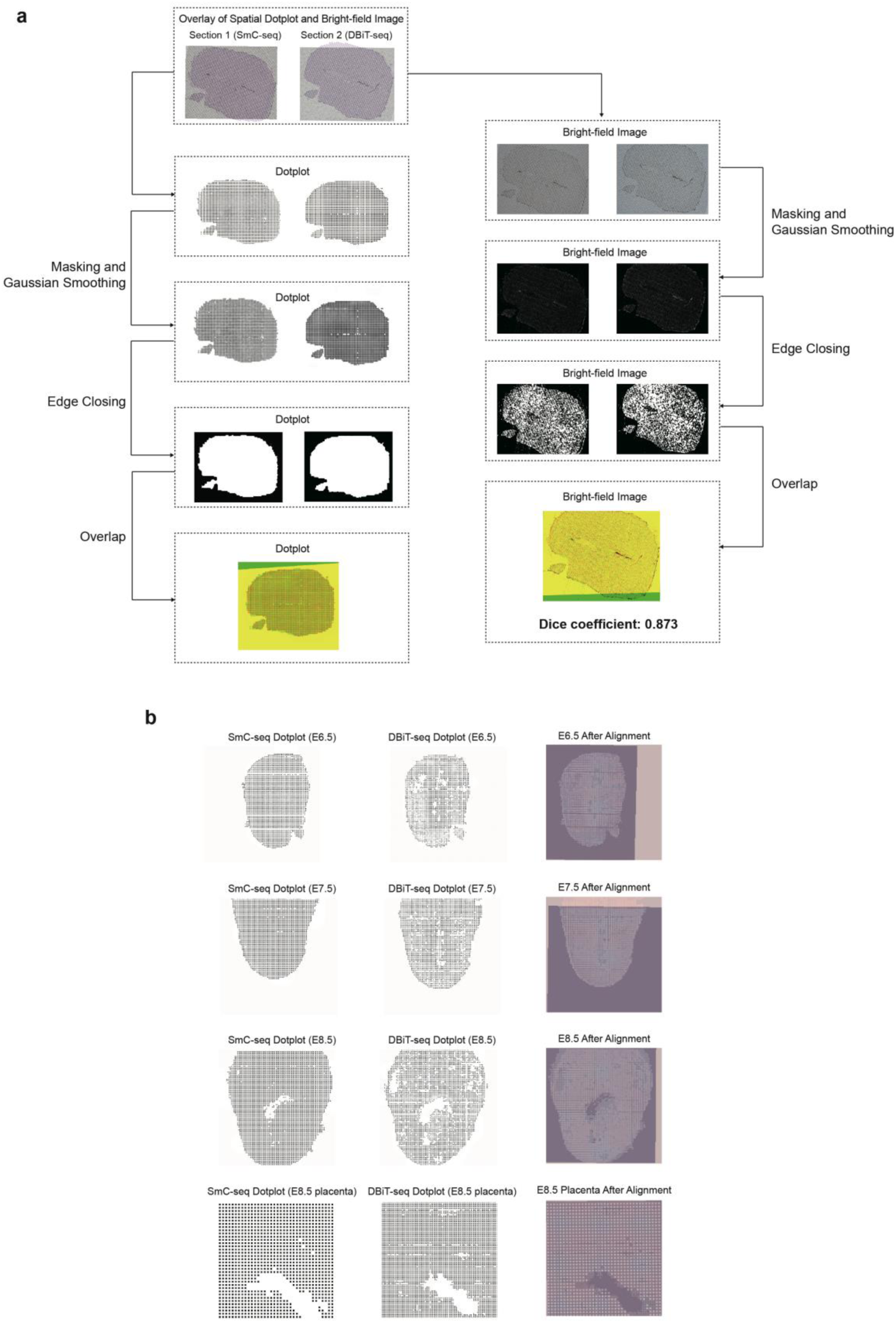
Tissue outline-based integration of SmC-seq and spatial transcriptomic sequencing. a, Workflow and validation of data integration based on tissue outline, illustrated using adjacent E6.5 tissue sections. The top panel shows an overlay of the dot plots and bright-field (BF) images, where each dot corresponds to a 10 μm-wide channel positioned over the tissue. The left panels illustrate the alignment workflow for dot plots using the tissue outline as reference. The right panel shows alignment of the BF image to the same outline, serving as validation. After alignment, the Dice coefficient was calculated based on the original BF image to quantify registration accuracy. **b,** Integration results of SmC-seq and DBiT-seq data from E6.5, E7.5, and E8.5 stages, aligned based on tissue outlines.

**Extended Data Fig.6.**
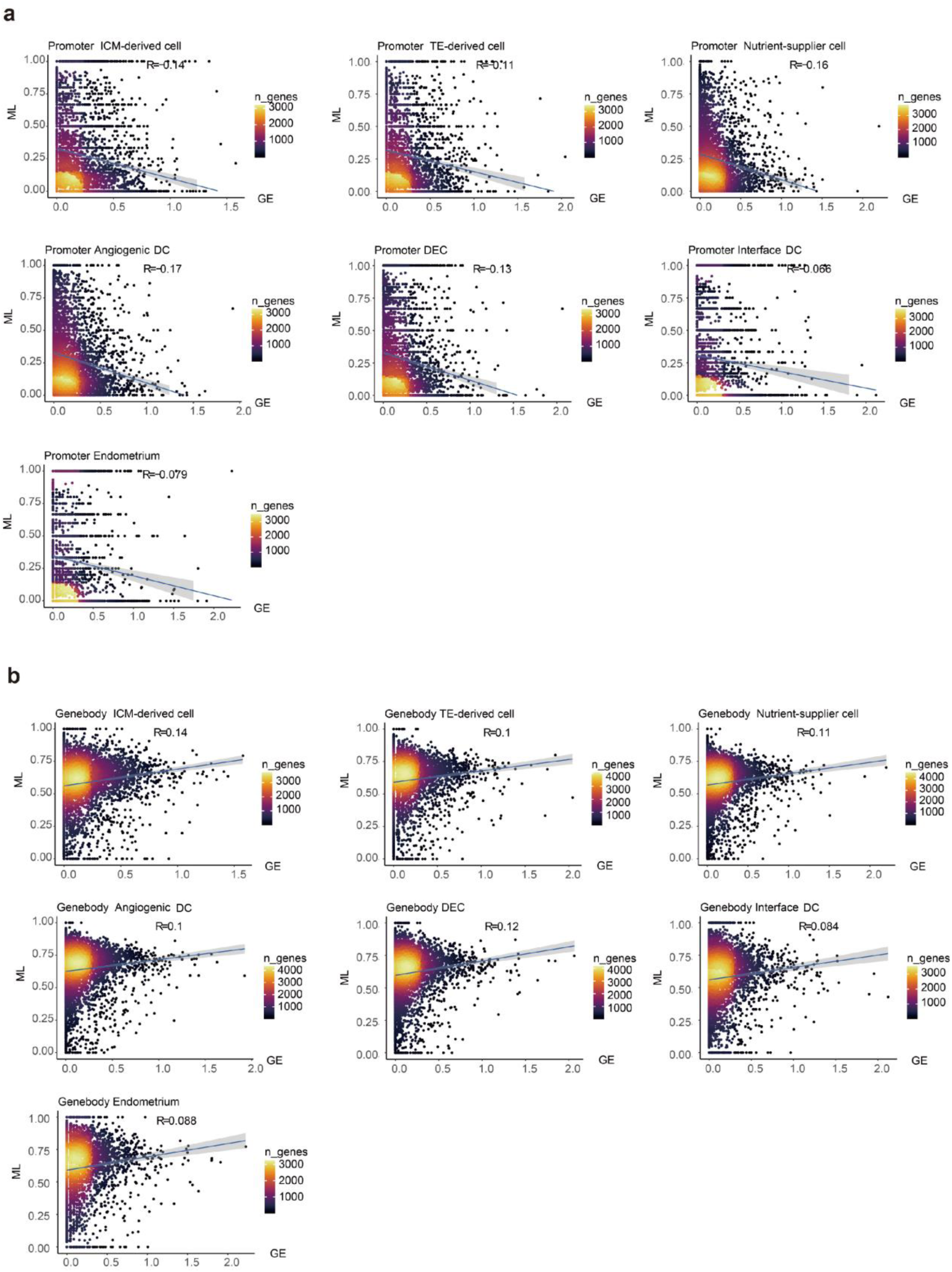
Global correlation between DNA methylation and gene expression across major clusters in mouse embryo-uterus tissue. a, Density plot showing the correlation between DNA methylation levels at promoter regions and corresponding gene expression levels for all genes across distinct clusters. The clusters include ICM-derived cell cluster, TE-derived cell cluster, nutrient-supplier cell cluster, angiogenic DC cluster, DEC cluster, interface DC cluster, and endometrium cluster. Linear regression lines are included. R, correlation coefficient; ML, methylation level; GE, gene expression; n_genes, number of genes. **b,** Density plot showing the correlation between genebody methylation levels and gene expression levels of all genes for the same clusters as in **a**. Linear regression lines are included.

**Extended Data Fig.7.**
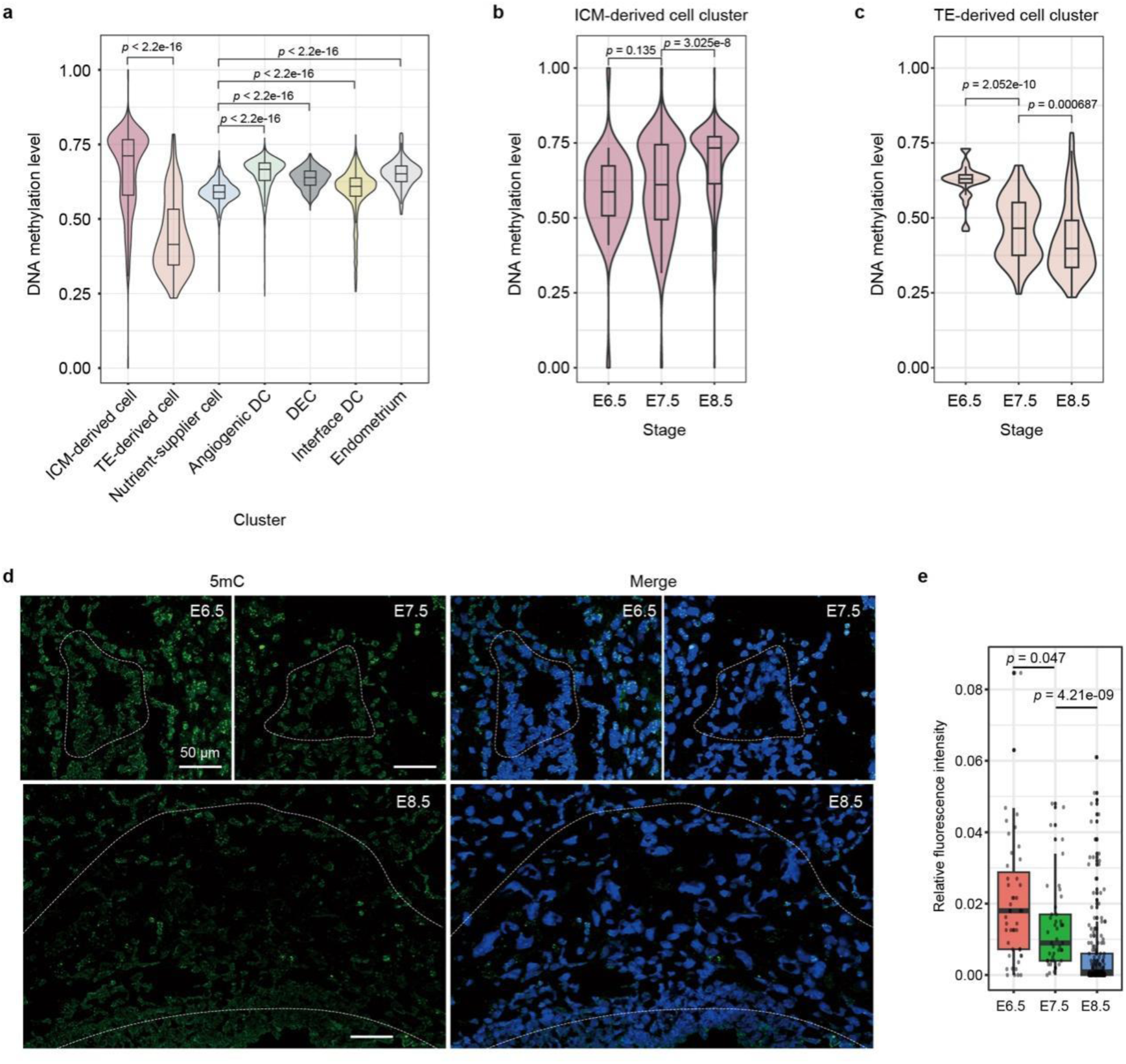
DNA methylation levels across major clusters in mouse embryo-uterus tissue after implantation. a, DNA methylation levels of pixels in different clusters within mouse maternal-embryo tissues after implantation. Two-sided wilcoxon rank sum tests were used. Boxes and whiskers represent the 25^th^/75^th^ percentiles and 1.5 x the interquartile range, respectively. ICM, inner cell mass; TE, trophectoderm; DC, decidual cell; DEC, decidual endometrial cell. **b-c**, Comparison of DNA methylation levels of pixels in ICM-derived cell cluster (**b**) and TE-derived cell cluster (**c**) among different developmental stages. Two-sided wilcoxon rank sum tests were used. **d,** Immunostaining for 5mC in EPC from the E6.5 to E8.5 stages. Dashed circles mark the EPC regions. Scale bar, 50 μm. Blue color represent DNA signal (Hoechst 33342). **e,** Statistical results of relative 5mC fluorescence signal from the E6.5 to E8.5 stages shown in panel **e**. Wilcoxon rank sum test was used for statistical test.

**Extended Data Fig.8.**
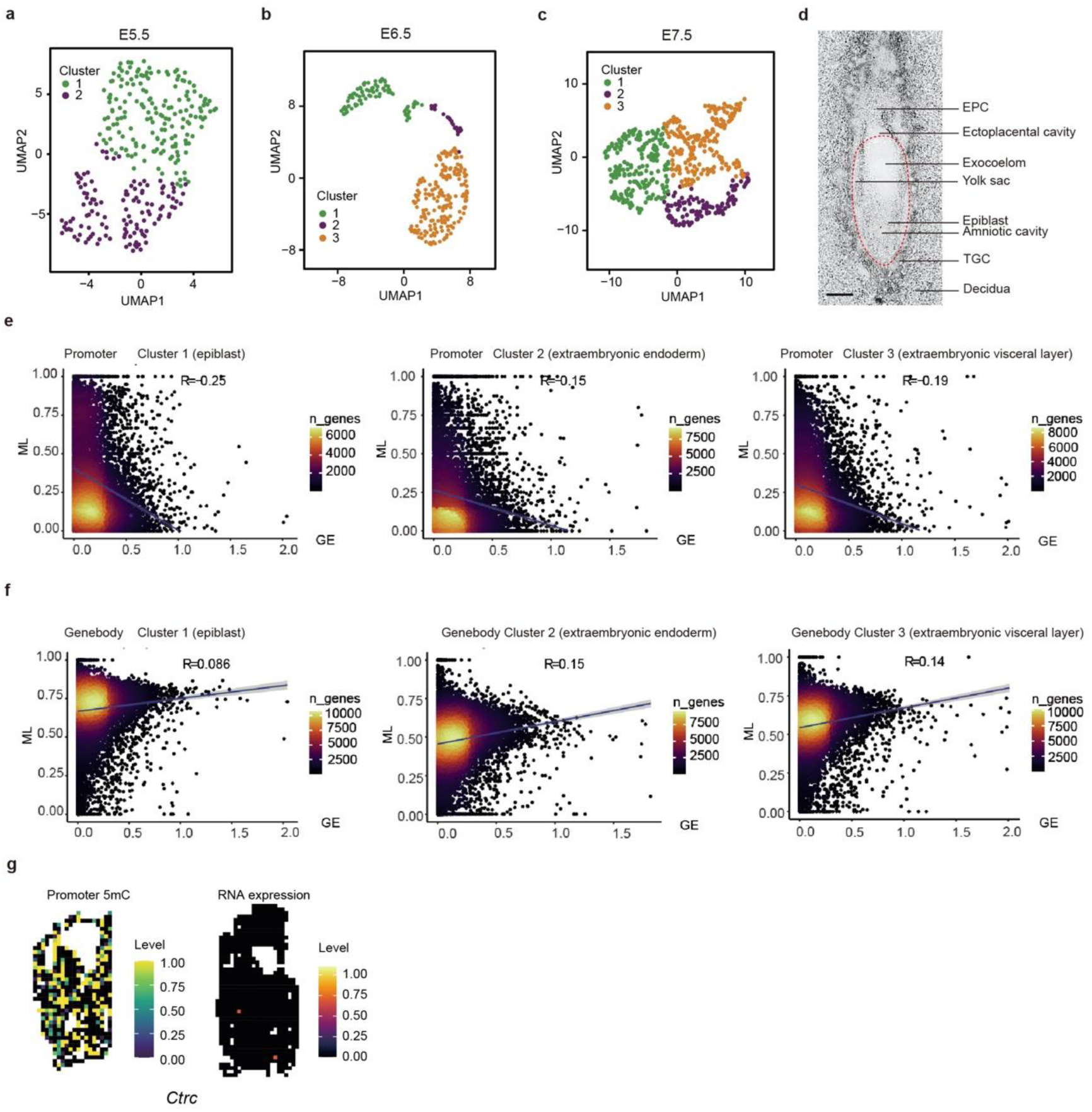
Clustering and methylation-expression correlation of ICM-derived cells. **a**, UMAP visualization of clusters derived from the E5.5 ICM, classified based on SmC-seq data. Fluidic chips with 10 μm-width microchannels were used to generate the data. **b**, UMAP visualization of clusters derived from the E6.5 ICM, classified based on SmC-seq data. Fluidic chips with 10 μm-width microchannels were used to generate the data. **c**, UMAP visualization of clusters derived from the E7.5 ICM, classified based on SmC-seq data. Fluidic chips with 10 μm-width microchannels were used to generate the data. **d**, Bright-field images of a frozen section of E7.5 embryo. The tissue outlined by the red dashed line is derived from ICM. Scale bar, 200 μm. **e**, Density plot showing the correlation between DNA methylation levels at promoter regions and corresponding gene expression levels for all genes across different clusters. R, correlation coefficient; ML, methylation level; GE, gene expression; n_genes, number of genes. **f**, Density plot showing the correlation between genebody methylation levels and gene expression levels for all genes across the same clusters as in panel **e**. g, Spatial distribution of promoter 5mC levels of *Ctrc* and its corresponding gene expression in an E7.5 embryo. Each pixel represents a 10 μm × 10 μm region. 5-μm-interval chips are used for this analysis.

**Extended Data Fig.9.**
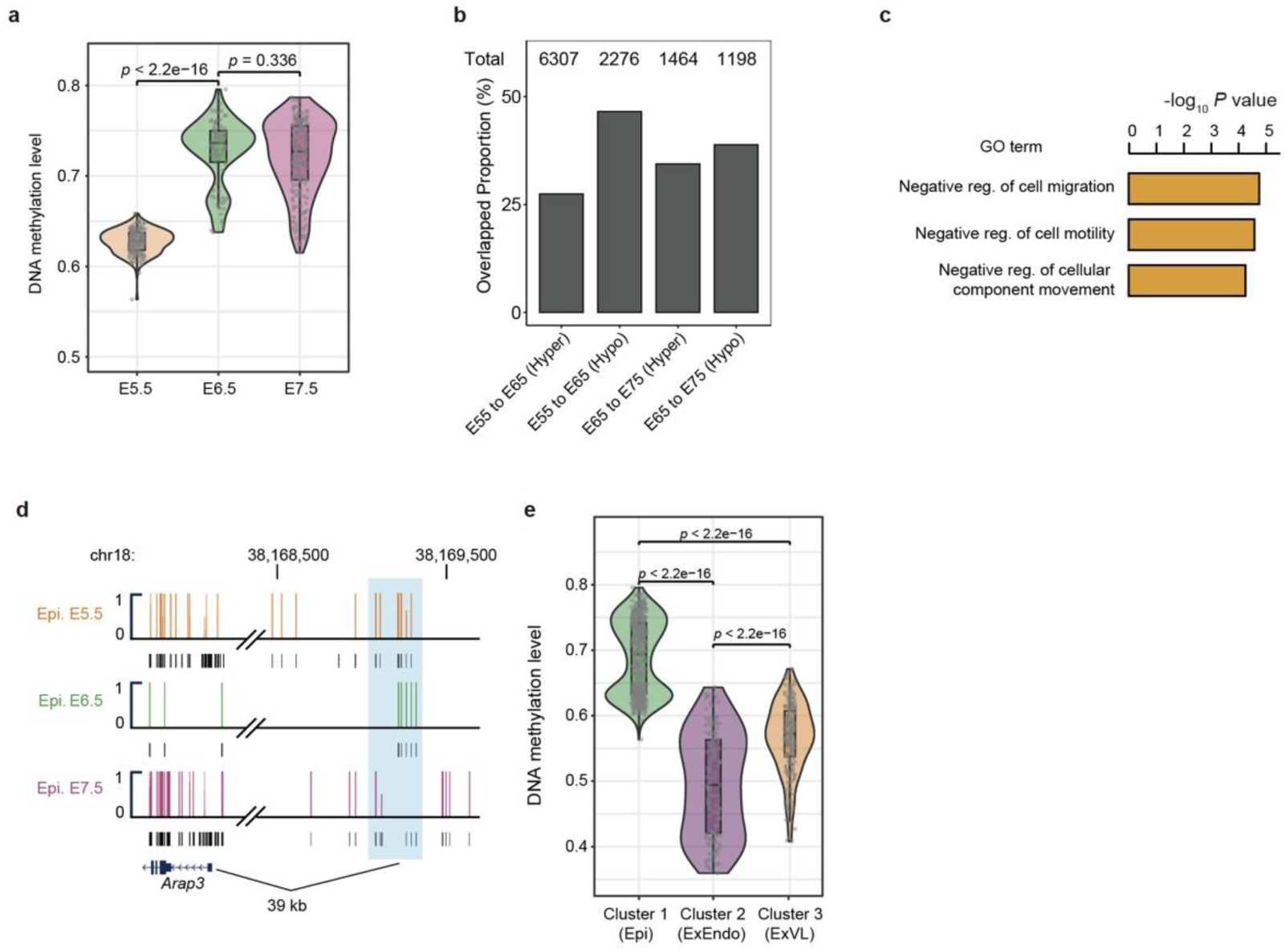
DNA methylation dynamics and functional analysis in epiblast cluster across developmental stages. a, Comparison of DNA methylation levels of cluster 1 (epiblast) at different developmental stages. Two-sided wilcoxon rank sum tests were used. Boxes and whiskers represent the 25th/75th percentiles and 1.5 x the interquartile range, respectively. **b,** Bar plot showing the percentages of hyperDMRs and hypoDMRs identified by SmC-seq that overlap with DMRs identified from published scNMT-seq data in mouse early embryo during gastrulation at the corresponding stages. The numbers of hyperDMRs and hypoDMRs detected by SmC-seq in epiblast cells from E5.5 to E7.5 stages are indicated above each bar. **c,** GO enrichment of genes associated with hypomethylated differentially methylated regions (hypoDMRs) in cluster 1 (epiblast) at the E7.5 stage compared to the E6.5 stage. reg., regulation. **d,** Genome browser view of DNA methylation levels of a hypoDMR in cluster 1 (epiblast) at the E7.5 stage that are associated with *Arap3*. Light blue shadow marks genomic region of the hypoDMR. Epi, epiblast. **e,** Comparison of DNA methylation levels of pixels from different clusters. Two-sided wilcoxon rank sum tests were used. Boxes and whiskers represent the 25th/75th percentiles and 1.5 x the interquartile range, respectively. ExEndo, extraembryonic endoderm; ExVL, extraembryonic visceral layer.

**Extended Data Fig.10.**
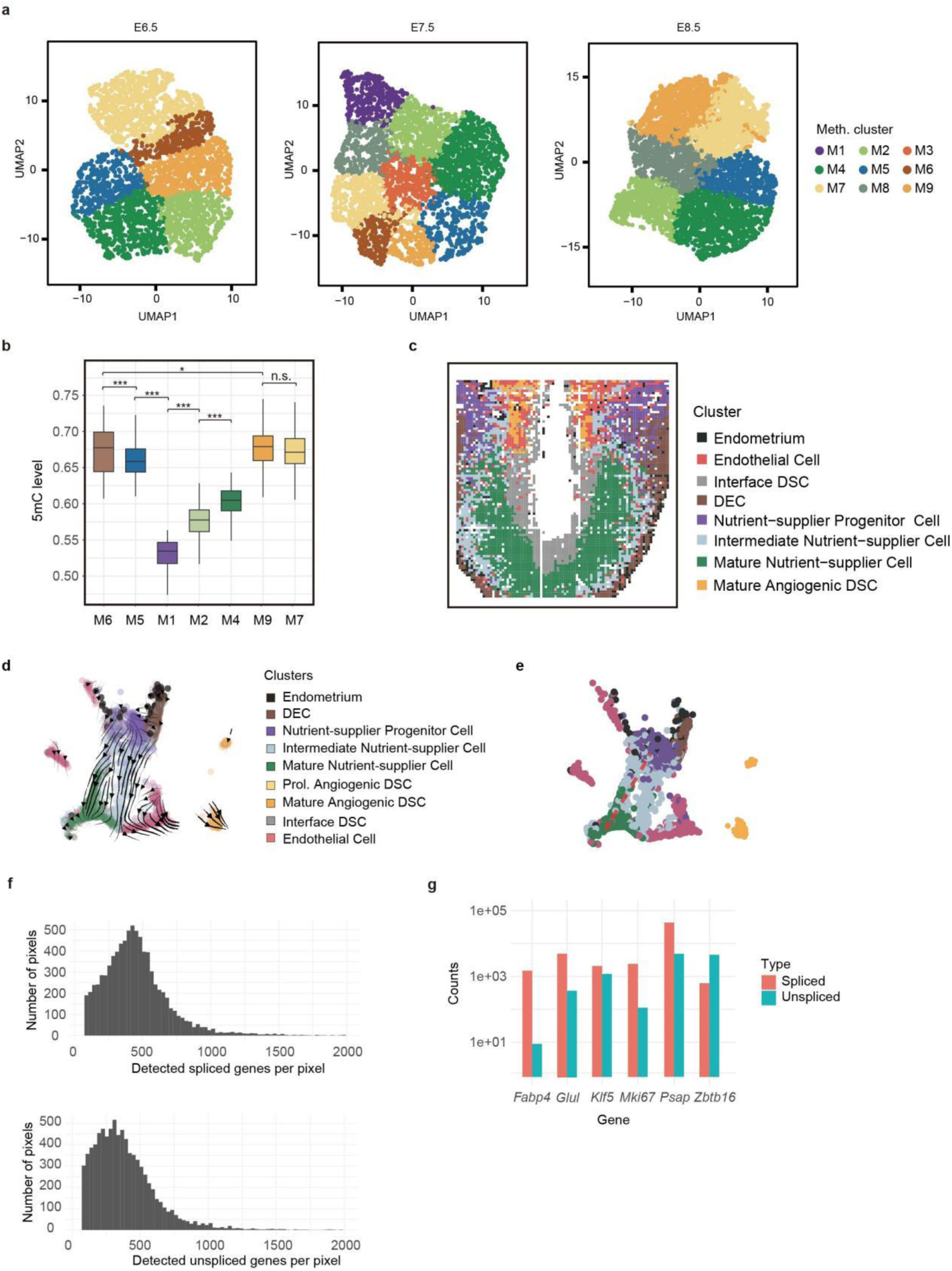
Clustering, methylation dynamics, and spatial distribution of maternal decidua. **a**, UMAP visualization of clusters in maternal decidual tissues at different developmental stages, classified based on SmC-seq data. **b**, Comparison of DNA methylation level among clusters classified based on SmC-seq data of maternal decidual cells. The global DNA methylation of pixels was calculated. Boxes and whiskers represent the 25th/75th percentiles and 1.5 x the interquartile range, respectively. Two-sided wilcoxon rank sum and signed rank tests were used. * represents p < 0.05; *** represents *p* value < 0.001; n.s. represents not significant. **c**, Spatial distribution of clusters in an E7.5 decidua at 10 μm × 10 μm resolution. **d**, RNA velocity analysis of spatial RNA-seq data from an E7.5 decidua section. Fluidic chips with 10 μm-width microchannels were used to generate the data. Each pixel represents a 10 μm × 10 μm region. **e**, UMAP visualization of spatial RNA-seq data from an E7.5 decidua section, with the inferred differentiation trajectory of nutrient-supplier cell lineage. The annotation of clusters is shown in panel d. **f,** Distribution of the number of detected spliced (top) and unspliced (bottom) genes per pixel in the dataset used for RNA velocity analysis. The x-axis shows the number of detected genes, and the y-axis shows the number of pixels. **g,** Total spliced and unspliced transcript counts across selected marker genes in the dataset used for RNA velocity analysis.

**Extended Data Fig.11.**
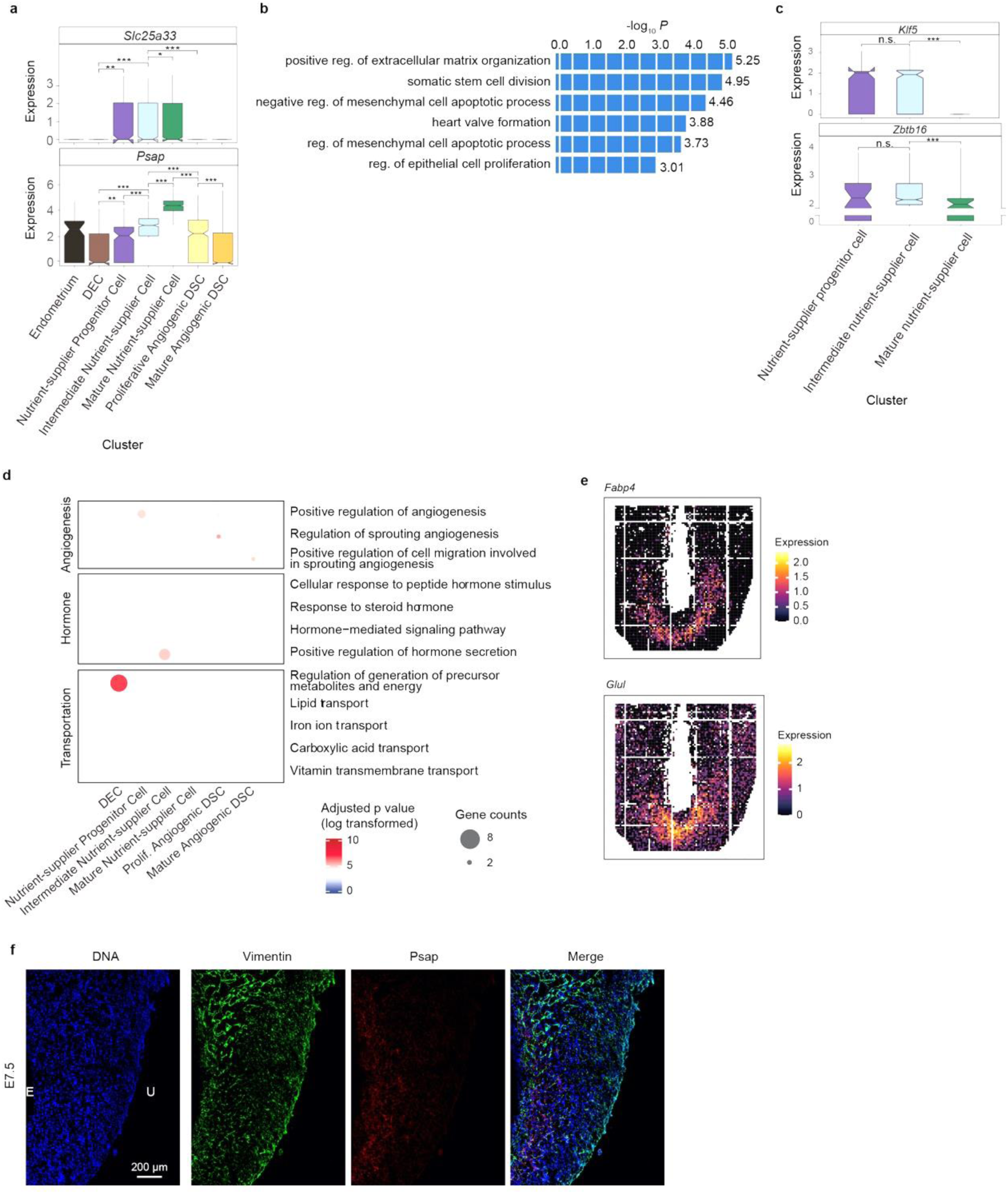
Expression and functional enrichment analysis in decidual cells. **a**, Boxplot showing the expression levels of *Slc25a33* and *Psap* in different maternal decidual clusters. Two-sided Wilcoxon rank-sum tests were used. * represents *p* < 0.05; ** represents *p* < 0.01; *** represents *p* < 0.001. **b,** GO analysis for the genes associated with the genomic regions (300 bp bins) that become hypermethylated in mature nutrient-supplier cell cluster compared to nutrient-supplier progenitor cell cluster. **c**, Boxplot showing the expression levels of *Klf5* and *Zbtb16* in different nutrient-supplier cell clusters. Two-sided Wilcoxon rank-sum tests were used. *** represents *p* value < 0.001; n.s. represents not significant. **d,** GO analysis of cell-type-specific marker genes across various maternal decidual clusters. The bubble size indicates the number of genes, while the color represents the significance. **e,** Spatial distribution of gene expression level of *Fabp4* and *Glul* in an E7.5 decidua. Each pixel represents a 10 μm × 10 μm region. **f**, Immunostaining of Vimentin and Psap in the lateral decidua at the E7.5 stage. Scale bar, 200 μm. E, embryo side; U, uterus side.

**Extended Data Fig.12.**
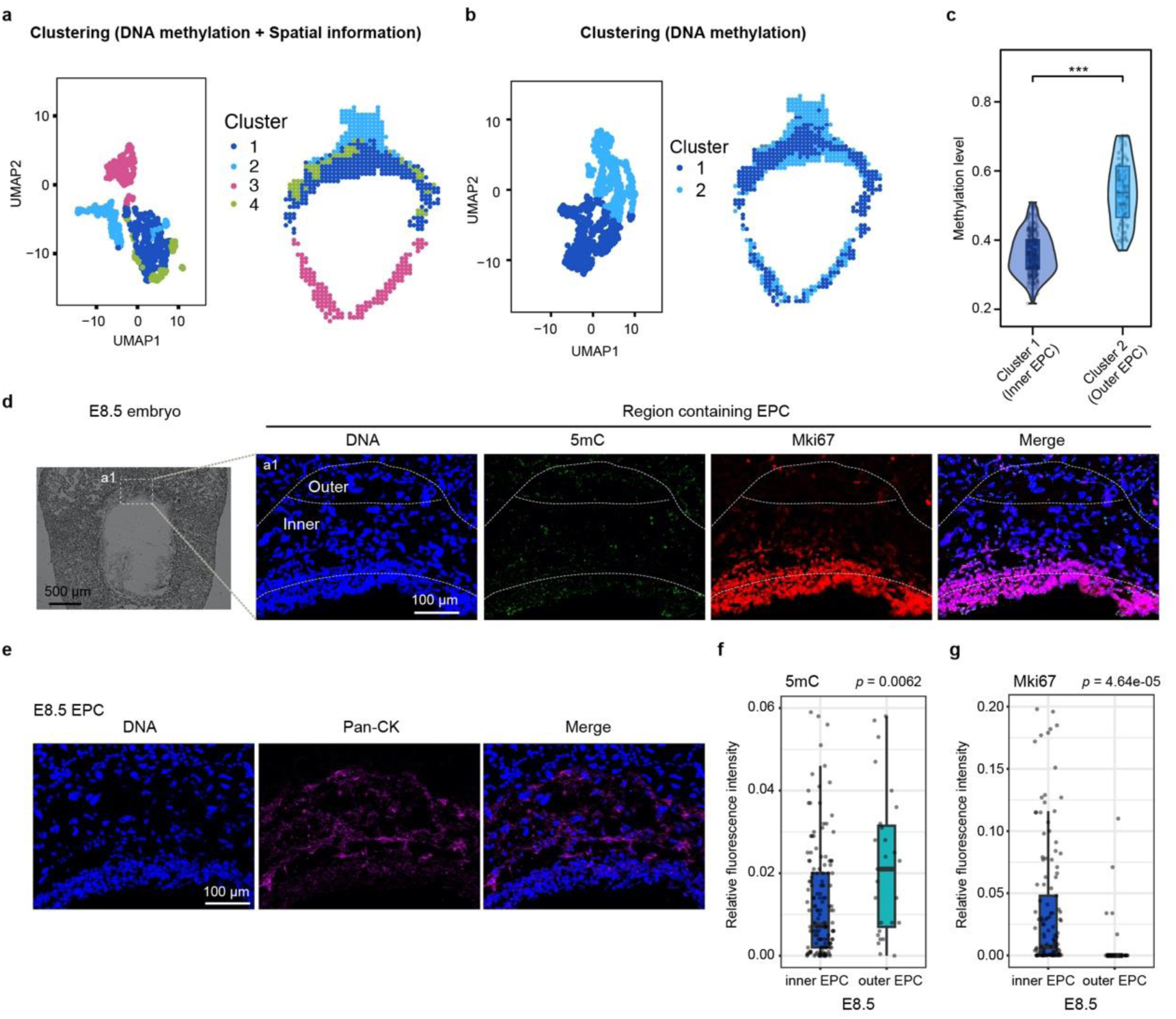
Clustering and comparison among TE-derived tissue using DNA methylation. a, UMAP visualization and spatial distribution of clustering for TE-derived tissue at the E8.5 stage, using both spatial distance matrix and DNA methylation data. **b**, UMAP visualization and spatial distribution of clustering for TE-derived tissue at the E8.5 stage, using DNA methylation data alone. **c,** Comparison of DNA methylation levels between clusters 1 and 2 based on SmC-seq data. The global DNA methylation level for each pixel was analyzed. Two-sided wilcoxon rank sum tests were used. *** represents *p* value < 0.001. Boxes and whiskers represent the 25^th^/75^th^ percentiles and 1.5 x the interquartile range, respectively. **d**, Immunostaining for 5mC and Mki67 in the ectoplacental cone (EPC) at the E8.5 stage. inner, inner EPC; outer, outer EPC. Dashed lines delineate inner and outer EPC regions. **e**, Immunostaining for pan-cytokeratin in the ectoplacental cone at the E8.5 stage. Pan-cytokeratin antibody was used to mark the EPC region for panel **d**. **f.** Statistical results of relative 5mC fluorescence signal between inner EPC and outer EPC shown in panel **d**. Two-sided wilcoxon rank sum test was used for statistical analysis. **g.** Statistical results of relative Mki67 fluorescence signal between inner EPC and outer EPC shown in panel **d**. Two-sided wilcoxon rank sum test was used for statistical analysis.

**Extended Data Fig.13.**
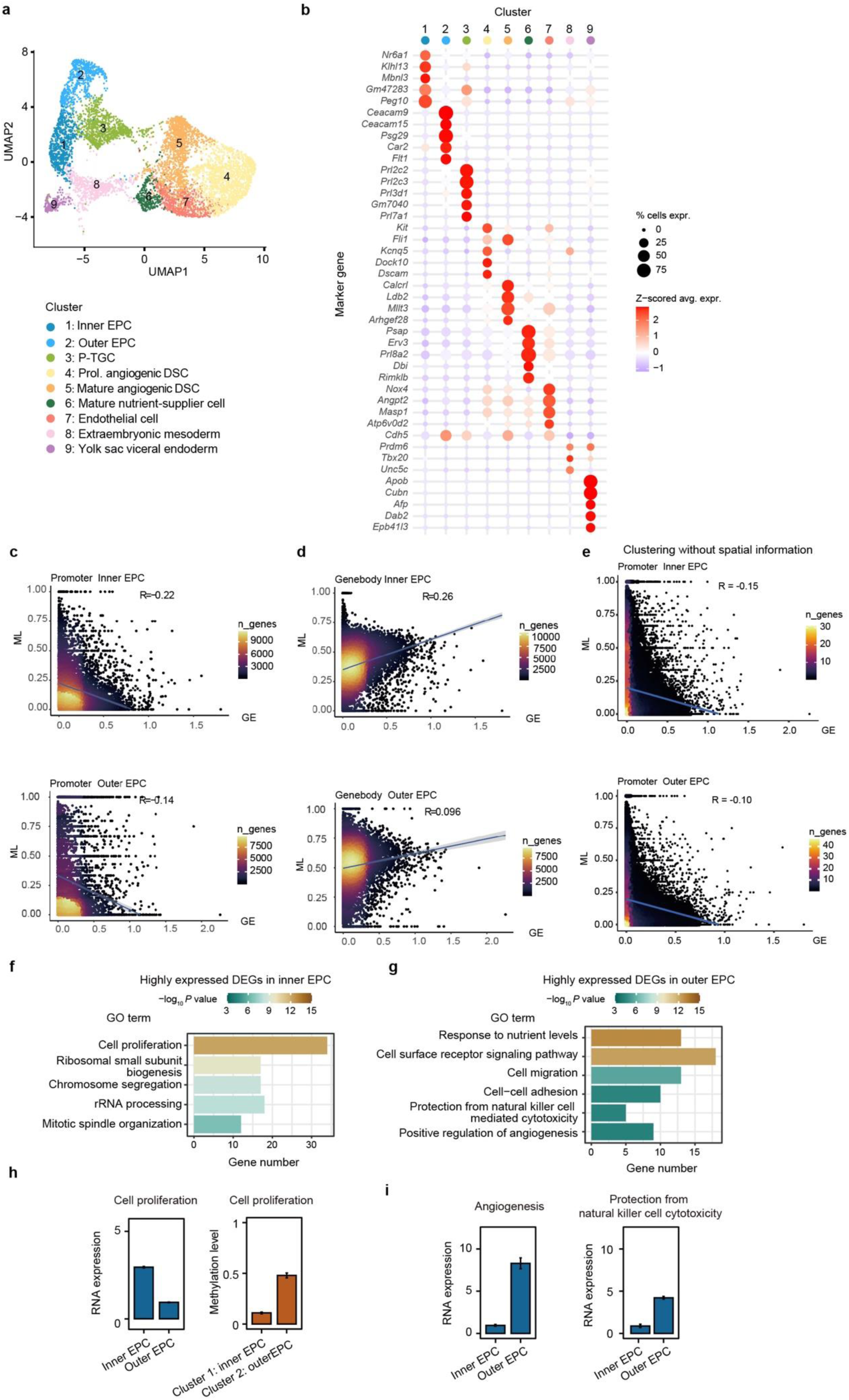
Clustering and comparison among TE-derived tissue using transcriptomic data. **a**, UMAP visualization of clusters classified based on spatial transcriptomic data of an E8.5 mouse tissue including TE-derived tissue. **b**, Heatmap of the top 5 marker genes for each cluster shown in panel **a**. The size of each point represents the percentage of pixels expressing that gene, the color indicates the average gene expression within each cluster. **c**, Density plot showing the correlation between DNA methylation levels at promoter regions and corresponding gene expression for all expressed genes across inner EPC and outer EPC. **d**, Density plot showing the correlation between gene body methylation levels and gene expression for the same clusters as in panel **c**. **e**, Density plot showing the correlation between promoter methylation levels and gene expression for the clusters identified without spatial information as shown in Extended Data Fig.12b. Cluster 1 represents inner EPC, cluster 2 represents outer EPC. **f**, GO enrichment of differentially expressed genes (DEGs) highly expressed in inner EPC. *P*-values were false discovery rate (FDR) adjusted. **g**, GO enrichment of differentially expressed genes (DEGs) highly expressed in outer EPC (right). *P*-values were FDR-adjusted. **h**, Average expression levels and promoter DNA methylation levels of genes associated with cell proliferation in inner EPC (cluster 1) and outer EPC (cluster 2). Error bars represent standard errors of the mean (SEM). **i**, Average expression levels of genes associated with angiogenesis (left) or protection from natural killer cell cytotoxicity (right) in inner and outer EPC clusters. Error bar represents SEM.

## Notes

### Competing Interest Statement

The authors have declared no competing interest.

